# The Generalizability of Cortical Area Parcellations Across Early Childhood

**DOI:** 10.1101/2024.09.09.612056

**Authors:** Jiaxin Cindy Tu, Michael Myers, Wei Li, Jiaqi Li, Xintian Wang, Donna Dierker, Trevor K. M. Day, Abraham Z. Snyder, Aidan Latham, Jeanette K. Kenley, Chloe M. Sobolewski, Yu Wang, Alyssa K. Labonte, Eric Feczko, Omid Kardan, Lucille A. Moore, Chad M. Sylvester, Damien A. Fair, Jed T. Elison, Barbara B. Warner, Deanna M. Barch, Cynthia E. Rogers, Joan L. Luby, Christopher D. Smyser, Evan M. Gordon, Timothy O. Laumann, Adam T. Eggebrecht, Muriah D. Wheelock

## Abstract

The cerebral cortex consists of distinct areas that develop through intrinsic embryonic patterning and postnatal experiences. Accurate parcellation of these areas in neuroimaging studies improves statistical power and cross-study comparability. Given significant brain changes in volume, microstructure, and connectivity during early life, we hypothesized that cortical areas in 1- to 3-year-olds would differ markedly from neonates and increasingly resemble adult patterns as development progresses.

Here, we parcellated the cerebral cortex into putative areas using local functional connectivity gradients in 92 toddlers at 2 years old. We demonstrate high reproducibility of these cortical regions across 1- to 3-year-olds in two independent datasets. The area boundaries in 1- to 3-year-olds were more similar to those in adults than those in neonates. While the age-specific group area parcellation better fit the underlying functional connectivity in individuals during the first 3 years, adult area parcellations might still have some utility in developmental studies, especially in children older than 6 years. Additionally, we provide connectivity-based community assignments of the parcels, showing fragmented anterior and posterior components based on the strongest connectivity, yet alignment with adult systems when weaker connectivity was included.

## Introduction

Understanding the intricate organization of the human brain is a fundamental pursuit in systems neuroscience. Previous research supports the notion that the cerebral cortex is divided into spatially contiguous areas distinguishable by function, architecture, connectivity, and/or topographic organization (Eickhoff et al., 2015; Felleman and Van Essen, 1991; Petersen et al., 2024). For example, the differentiation between prestriate and striate areas by a clear histochemical border has been observed in human and monkey fetuses (Kostovic and Rakic, 1984). Besides the historical approaches in using histology to find the area organizations(Amunts et al., 2013; Amunts and Zilles, 2015; Brodmann, 1905; Carmichael and Price, 1994; Evans, 1992), connectivity-based area parcellations, especially those with fMRI data, has become popular as an efficient and non-invasive alternative to parcellate brain areas (Eickhoff et al., 2018, 2015; Glasser et al., 2016; Gordon et al., 2016; Schaefer et al., 2018; Shen et al., 2013). These connectivity-based area parcellations relies on the computation of connectivity strength to other parts of the brain for each voxel/vertex (a.k.a. connectivity profiles) and then group the voxel/vertex into areas with homogeneous connectivity profiles (Eickhoff et al., 2015).

The formation of cortical areas occurs starting from embryonic development. Initially, continuous gradients of signaling molecules within the ventricular zone drive the formation of neurons from their progenitor cells and give rise to a “protomap” (Bishop et al., 2002; Cadwell et al., 2019; Fukuchi-Shimogori and Grove, 2001; Hamasaki et al., 2004; O’Leary et al., 2007; Rakic, 1988; Stiles and Jernigan, 2010). Later, both intrinsic and extrinsic factors refine this “protomap” into discrete areas (Cadwell et al., 2019; O’Leary et al., 2007; Qian et al., 2024). One important contributor to this process is environmental inputs (Catalano and Shatz, 1998; Greenough et al., 1987; Smyser et al., 2011; Tau and Peterson, 2010), especially from the thalamocortical axon projections (Molnár and Kwan, 2024; O’Leary et al., 2007; Vue et al., 2013). For example, early deprivation of vision in one eye caused shifted ocular dominance columns in monkeys (Hubel et al., 1997). The explosive increase in exposure to environmental stimuli following birth likely plays a significant role in the refinement of area boundaries shortly after birth. Moreover, synaptic addition and growth of dendrites and spines also enters a phase of logarithmic growth in the first few months after birth (Levitt, 2003), suggestive of an elevated period of cortical plasticity. Considering these factors, it is reasonable to expect that cortical areas in neonates would show low similarity to those in adults (Myers et al., 2024), with greater similarity to adult brain areas as the brain develops. Furthermore, it has been postulated that developmental changes are not uniform across the brain. The sequence of development has previously been described to follow a sensorimotor-to-association axis (Casey et al., 2005; Dean et al., 2015; Flechsig, 1901; Grayson and Fair, 2017; Hill et al., 2010; Smyser and Neil, 2015; Smyser et al., 2016; Sydnor et al., 2021; Tau and Peterson, 2010; von Economo et al., 2008; von Economo and Koskinas, 1925), or a posterior-to-anterior axis (Larivière et al., 2020; Q. Li et al., 2024). Few studies have examined whether the maturation of cortical areas followed either of these patterns.

Many neuroimaging analyses have been conducted at the scale of parcels (Arslan et al., 2018; Bijsterbosch et al., 2020; Farahani et al., 2019; Faskowitz et al., 2022, 2022; Helwegen et al., 2023; Luppi et al., 2024; Zalesky et al., 2010). Inaccurate area parcellation choice can lead to the mixing of signals (Smith et al., 2011), conceal known community structure (Power et al., 2011), and reduce the prediction accuracy of clinical phenotypes (Abraham et al., 2017). Therefore, choosing an area parcellation scheme that closely reflects the actual area boundaries in the data is of great importance for functional connectivity (FC) analyses (Grayson and Fair, 2017).

Neuroimaging analyses often adopt definitions of cortical areas in adult brains (Glasser et al., 2016; Gordon et al., 2016; Schaefer et al., 2018; Shen et al., 2013). However, the dynamic and rapid development of the brain during infancy (Bethlehem et al., 2022) triggers unique concerns about whether it is valid to apply existing adult area parcellations to early childhood brains (Cusack et al., 2018; Oishi et al., 2019; Shi et al., 2018; Wang et al., 2023). In response, several early childhood area parcellations have been developed in recent years (Myers et al., 2024; Scheinost et al., 2016; Shi et al., 2018; Wang et al., 2023). Despite these advances, having different area parcellations for different age ranges poses a practical challenge for making coherent comparisons in brain organization across development. Thus, many researchers have continued to use adult area parcellations in early childhood studies (Kim et al., 2023; Nielsen et al., 2022; Yates et al., 2023), as well as studies across the lifespan (Betzel et al., 2014; Cao et al., 2014; Puxeddu et al., 2020; Zuo et al., 2017).

One crucial factor in determining which area parcellation to employ in a given age range would be the degree to which an age-specific area parcellation differs from an adult area parcellation in pediatric samples. However, a systematic examination of parcellations across age groups is lacking. We aim to a) illustrate how well the area parcellations fit the functional connectivity data across individuals at various developmental stages, b) quantify the improvement compared to adult parcellations, and c) evaluate the potential impact of using an adult parcellation instead of the proper early childhood parcellation on downstream analyses. If adult parcellations separate the cortical areas with comparable success as early childhood parcellations, utilizing adult parcellation schemes for developmental cohorts would be justifiable. One prior study suggested that this was not the case for neonates (Myers et al., 2024). Here we query whether the adult parcellation would be a reasonable choice for older infants, toddlers and children.

In the current study, we derive a surface-based area parcellation based on FC local gradient transitions (Cohen et al., 2008; Gordon et al., 2016; Wig et al., 2014) in 92 toddlers at age of 2 years. To test the reproducibility of our area parcels across groups of subjects and whether the reproducibility followed a uniform distribution across space, we derive parcellations using half the sample (n = 46). To examine differences in patterns of FC local gradient transition across development, we quantify the similarity between the boundary maps at different developmental stages. Furthermore, we compare our area parcellation to alternative adult and early childhood parcellations and demonstrate the generalizability and limitations of our area parcellation for application to various developmental stages. Finally, we derive the community organization which describes the relationship between the area parcels.

## Methods

Experiments were undertaken with the understanding and written consent of each subject or their parents for all datasets used in this project.

### Neuroimaging Data for Deriving Area Parcellations

One main goal of this paper is to examine the area parcellations at ages 1-3. We used two early childhood datasets: eLABE (Y2) and BCP (Table 1). The early childhood datasets used in the current study were all collected with a Siemens Prisma 3T scanner using HCP-style acquisition parameters (Supplementary Table 1). The functional MRI acquisition lasts 420 frames per scan run with 2-4 runs in the Baby Connectome Project (BCP) and 1-8 runs in the Early Life Adversity, Biological Embedding (eLABE) 2-year-old data (Y2). Anatomical scan processing and segmentation were conducted using age-specific pipelines (Kaplan et al., 2022). Functional data preprocessing followed established procedures (Power et al., 2014). Toddler EPI BOLD preprocessing pipeline was used for eLABE (Y2) and DCAN-Infant v0.0.9 (Autio et al., 2020; Donahue et al., 2016; Glasser et al., 2013) were used for BCP. Motion correction was performed with rigid-body transforms. The functional data were also corrected for asynchronous slice time shifts and systematic odd-even slice intensity differences attributable to interleaved acquisition (Power et al., 2012). The data were intensity normalized to achieve a consistent whole-brain mode value, and subsequently resampled to atlas space before being projected onto the 32k_fs_LR standard surface (Van Essen et al., 2012). Denoising was accomplished by nuisance regression, with regressors consisting of a 24-parameter Volterra expansion of motion time series, the mean signal over gray-ordinates, and the mean signals derived from white matter and cerebrospinal fluid (CSF) compartments. The data were bandpass filtered to retain BOLD-specific frequencies and geodesically smoothed with Connectome Workbench (Glasser et al., 2013; Marcus et al., 2011). Frame censoring was performed based on the frame displacement time series (FD > 0.2mm) following age-specific notch-filtering to exclude respiratory frequencies (Kaplan et al., 2022). Structural and functional scans were manually inspected and runs/sessions that failed quality controls were discarded. Additionally, participants who were born preterm (<37 weeks gestational age), had any neonatal ICU experience, or had signs of injury on MRI were also excluded from the analysis. Functional data with less than 600 low-motion frames were also excluded. For additional dataset-specific details, see Supplementary Table 1.

**Table 1.**
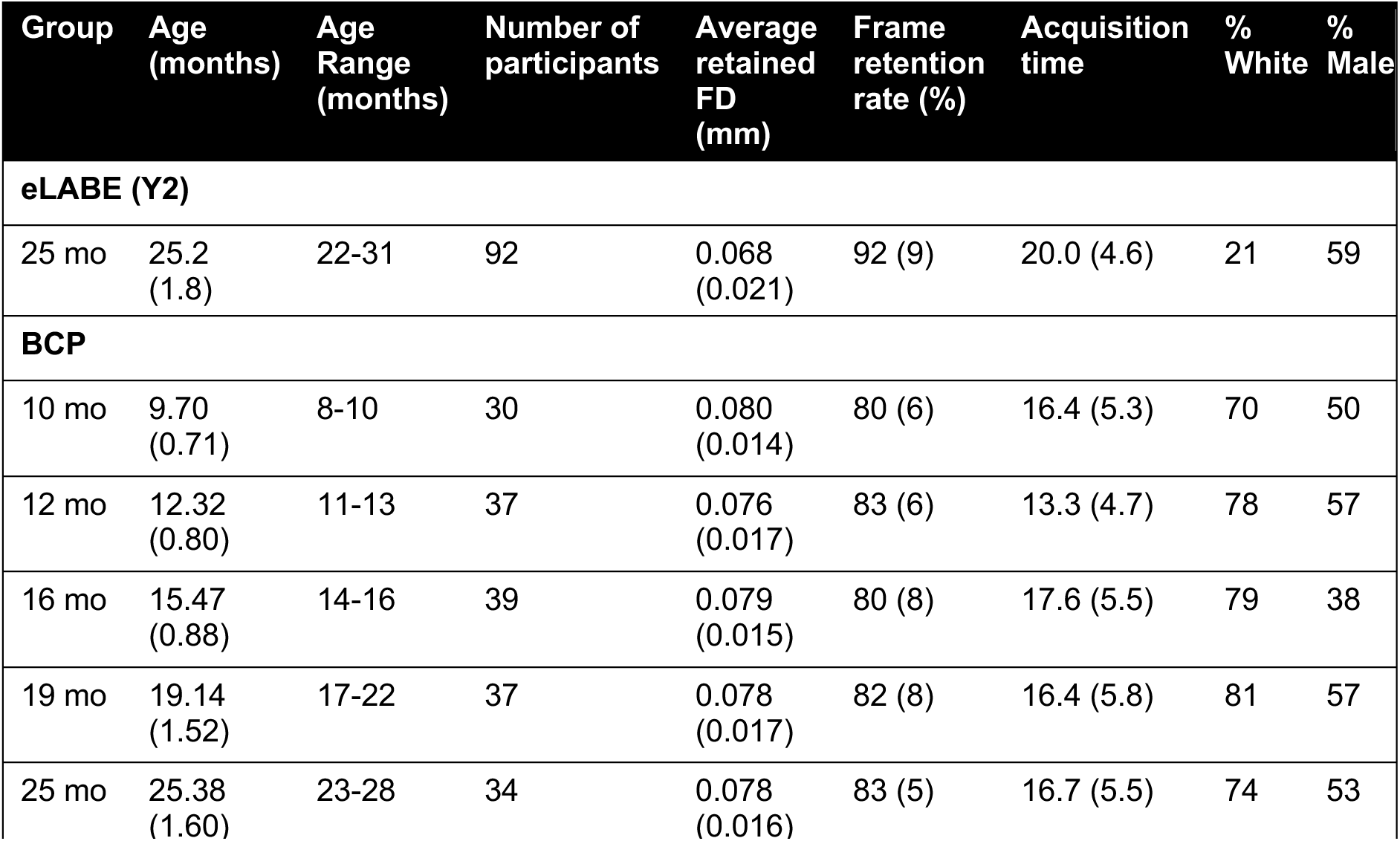
Subject demographics for the two infant/toddler datasets. For continuous variables, the mean is provided along with standard deviations in brackets. The group identity was defined as the median age rounded to the nearest whole number.

A summary of the demographics and image quality of the developmental cohort discovery and validation datasets is provided in Table 1. The cross-sectional age distribution and the distribution of age in longitudinal sessions are displayed in Supplementary Figure 1.

### Neuroimaging Data for Comparing FC Boundaries Across the Lifespan

To compare FC boundaries, we additionally included the FC boundaries from a young adult dataset (Washington University 120, WU120) used in a widely adopted adult parcellation (Gordon et al., 2016) and from the same neonate dataset (eLABE (Birth)) used in a neonatal parcellation (Myers et al., 2024). Acquisition and processing of these datasets followed similar pipelines to the early childhood datasets above and as briefly described below. For dataset-specific details, please refer to Supplementary Table 2.

#### WU120

Data were collected from 120 healthy young adult participants recruited from the Washington University community during relaxed eyes-open fixation (50% male, ages 19–32). Scanning was conducted using a Siemens TRIO 3T scanner and included the collection of high-resolution T1-weighted and T2-weighted images, as well as an average of 14 min of resting-state fMRI. Detailed acquisition and processing have been reported previously (Power et al., 2014).

#### eLABE (Birth)

Inclusion criteria were the same as the eLABE (Y2) cohort. Neuroimaging data were collected in 261 full-term, healthy neonate offspring shortly after birth (average postmenstrual age of included participants 41.7 weeks, range 39–45 weeks, 54% male). A total of 131 participants with the most data following frame censoring were used to create the FC boundaries. Additional details are in Supplementary Table 2.

### Neuroimaging Data for Testing the Generalizability of Areas Across the Lifespan

To test for the generalizability of area parcellations across the lifespan, we additionally include the year-3 timepoint from the eLABE dataset, Healthy Brain Network (HBN) children dataset, and HCP young adult (HCP-YA) dataset.

#### eLABE (Birth)

This is the same dataset as above. Because 131 of the participants were involved in the creation of the Myers-Labonte parcellation (Myers et al., 2024), the other 130 participants *not used in the generation of Myers-Labonte parcellation* were used to test the parcellation’s cluster validity performance to prevent circularity. The acquisition protocol and processing pipeline were the same as described before (Supplementary Table 2).

#### eLABE (Y3)

The inclusion criteria were the same as the eLABE (Y2) cohort. Neuroimaging data were collected from 132 participants at the age of 3 years. Additional participants were excluded based on the quality of structural and functional data and having less than 8 min (600 frames) of low-motion (respiratory-filtered FD < 0.2) data retained, leaving 65 participants (range = 2.93-3.97 years, mean = 3.22 years, SD = 0.32 years, 63% male). The acquisition protocol and processing pipeline were the same as the eLABE (Y2) dataset at age two.

#### HBN

Resting-state fMRI data from 493 participants from the first nine releases of the Healthy Brain Network (HBN) were divided into 10 groups by year (6-15yr). The <u>HBN study</u> is a large, multi-site study of children and young adults ages 5–21 years all collected in the New York area. Recruitment, consent, and study procedures are described in the data publication (Alexander et al., 2017) as well as project <u>website</u>. We used the data from two sites (CitiGroup Cornell Brain Imaging Center (CBIC) and Rutgers University Brain Imaging Center (RUBIC)).

Data were pre-processed using the Human Connectome Project minimal processing pipeline (Glasser et al., 2013). Additional processing steps (demeaning, detrending, nuisance regression (with regressors consist of a 24-parameter Volterra expansion of motion time series, the mean signal over gray-ordinates), bandpass filtering at 0.008-0.1 Hz to retain BOLD-relevant frequency and frame censoring at respiratory-filtered FD > 0.2 mm were carried out using custom-written Python (v3.8) scripts using the numpy v1.24.4, scipy v1.10.1, nibabel v5.1.0, and pandas v2.0.3 libraries. Each scan session takes 10 min and all included sessions comprises at least 8 min (600 frames) of low-motion (respiratory-filtered FD < 0.2) data retained. Data was geodesically smoothed to achieve an effective smoothing of 2.55 sigma gaussian kernel.

#### HCP-YA

Resting-state fMRI data from a subset of randomly chosen 40 participants not used for the creation of the Glasser parcellation (Glasser et al., 2016) were selected from the HCP-YA dataset for external validation of the adult dataset to minimize circularity. Data was processed with the same standard preprocessing pipeline as WU 120, except that a low-pass-filtered FD < 0.04 mm was used to remove high-motion frames.

### Creation of FC-transition Boundary Maps and Area Parcels

We segmented the cortical surface into discrete parcels representing putative cortical areas based on the FC local gradient (Gordon et al., 2016). The FC from each vertex to every other vertex was calculated as Pearson’s correlation of the time series in individual sessions (Supplementary Figure 2A). The Fisher-Z-transformed FC from each vertex was correlated with a randomly subsampled set of 594 vertices (1% of the total vertices) to generate an “FC similarity” matrix, which indexed the similarity in FC patterns across vertices (Supplementary Figure 2B). We used 1% of the vertices for computational efficiency without compromising accuracy (Supplementary Materials). After that, the workbench command “cifti-gradient” was used to calculate the gradient of FC-transition in individual subjects’ surfaces in the standard 32k_fs_LR mesh. The gradient maps were then averaged across all subjects and smoothed with a Gaussian kernel of 2.55 sigma (Supplementary Figure 2C). A “watershed by flooding” algorithm (Beucher and Meyer, 1992) was used to create discrete areas separated by boundaries based on the gradient transitions (Supplementary Figure 2D). The gradient-based boundary map technique rests on the assumption that FC within a cortical area is relatively uniform and distinct from FC of an adjacent area (Wig et al., 2014), consistent with how areas have been previously reported as distinct in connectivity in macaque monkeys (Felleman and Van Essen, 1991). Finally, the boundaries from different gradient maps were averaged to obtain a boundary map that indexed the probability of a vertex being an area boundary (values range between 0 and 1) (Supplementary Figure 2E).

Discrete parcels (Supplementary Figure 2F) from a boundary map were created by locating the minima in the boundary map, growing parcels from minima using the watershed algorithm, and merging the watersheds if the median values of boundaries between them are below a threshold (merging threshold, defined as a percentile of the boundary map values). Neighboring parcels with sizes smaller than 15 vertices were merged. Parcels joined only by a single vertex were split. Isolated parcels smaller than 10 vertices were removed. Vertices above 90% in the boundary map values (height threshold) were left as parcel borders. The resolution of the parcels depends on the merging threshold, with higher merging thresholds leading to a small number of larger parcels and lower merging thresholds leading to a large number of smaller parcels (as demonstrated in Supplementary Figure 3).

### Parcel Reproducibility

To assess the reproducibility of our results, we generated the boundary map (Supplementary Figure 2E) and discrete parcels (Supplementary Figure 2F) from non-overlapping split halves of participants 20 times. For each pair of area parcellations at a merging threshold, we quantified the overlap in the parcels and in the boundaries (See Section: **Parcel Similarity Measures**). In addition, we divided the brain into 10 equal bins based on either the position along the sensorimotor-association axis (Sydnor et al., 2021) or the posterior-anterior axis and calculated the parcel reproducibility in each bin (Supplementary Figure 4).

### Parcel Similarity Measures

Adjusted Rand Index (ARI) calculated on non-boundary vertices was used as the main measure of similarity across two area parcellations. For completeness and comparability with prior literature, we also calculated the dice coefficient between area parcellations, either as the average dice coefficient across matching pairs of parcels defined with the largest dice coefficient (Shen et al., 2013), or on binarized parcel identity maps (with boundaries as 0 and parcels as 1) (Myers et al., 2024). The dice coefficient between binarized parcel identity maps was biased by the percentage of the brain covered with parcels (e.g. when there are more/wider boundaries, the overlap will be higher). For example, in area parcellations such as Glasser (Glasser et al., 2013) and AAL (Tzourio-Mazoyer et al., 2002), the dice coefficient calculated this way would be 1 because those area parcellations did not specify boundaries and allocate all cortical vertices into parcels.

Additionally, we compared the binarized parcel boundaries (with boundaries as 1 and parcels as 0). We quantified the differences between the parcel boundaries with dice coefficient and Hausdorff distance (Müller et al., 2022; Shen et al., 2013), which measures the maximum distance one needs to travel between two contours. A lower Hausdorff distance indicates a high similarity between boundaries. We used a spatial distance measure for boundaries because it is less sensitive to small shifts in space and does not require perfect overlap. To mitigate sensitivity to outliers, we used two variants of the Hausdorff distance measure: 95% Hausdorff distance (HD95) (Huttenlocher et al., 1993) and average Hausdorff distance (AHD) (Müller et al., 2022). HD95 was defined as the maximum of the 95^th^ percentile of the distances between any point in contour X to the closest point in another contour Y and the 95^th^ percentile of the distances between contour Y to the closest point in contour X. AHD was defined as the maximum of the mean distance between contour X and contour Y and the mean distance between contour Y and contour X.

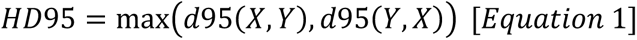

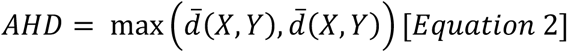

Here “distance” was defined as the geodesic distance between vertices in the Conte69 surface atlas (Van Essen et al., 2012).

Additionally, we generated a null model by generating random rotations around the x,y, and z axes for a split-half of the total sample of data (Split-I) and calculated each of the metrics. In theory, this controls for the bias from different merging thresholds, but due to the presence of the medial wall, spatial permutations often induce missing data (Markello and Misic, 2021).

### Boundary Map Consistency Across Age

To examine the difference in the area organization across different developmental stages, we applied the same method to generate boundary maps from neonates and adults. These were compared to the boundary maps derived from the eLABE (Y2) dataset. We computed the similarity between the boundary maps by taking the top percentiles of the boundary map values and calculating the Hausdorff distance measures.

### Evaluation of Cluster Validity of Area Parcellation

To evaluate the cluster validity of the area parcellations (i.e. how well they fit the FC data, we used an unbiased metric for the comparison of area parcellations across different spatial resolutions (Zhi et al., 2022). The distance-dependent boundary coefficient (DCBC) (Zhi et al., 2022) compares the average difference in similarity (Pearson’s r, with a value between −1 and 1) of FC profiles from vertices within a parcel and those from vertices between parcels across geodesic distance bins of 1 mm (e.g. between 10 mm to 11 mm). As demonstrated in a prior publication (Zhi et al., 2022), this metric accounts for the spatial smoothness of the data and is relatively unbiased when comparing area parcellations across multiple spatial resolutions (a.k.a. number of parcels). The expected value of DCBC for a random parcellation was zero regardless of the resolution of the parcellation, and a positive DCBC would mean better than random. Thus, no simulation with random null parcellations is necessary to establish a baseline measurement, as opposed to measures like homogeneity Z-score compared to a spatially permuted null (Gordon et al., 2016). As a negative control, we also evaluated an area parcellation that randomly partitioned the brain into 304 equally-sized fragments (Icosahedron) as a control. For implementation details and a comparison with alternative measures, please refer to the Supplementary Materials.

### Comparing Our Area Parcellation to Alternatives

To further contextualize results, we compared our area parcellation to existing area parcellations created using adult or early childhood data. We transformed the area parcellations into the common 32k_fs_LR standard mesh where necessary. Details for the transformation are provided in the Supplementary Materials.

Table 2 summarizes the area parcellations tested including the number of parcels, sources, and original space. In addition, to establish a lower bound of DCBC for the dataset, we used an Icosahedron-162 parcellation which provided regular tessellations of the hemispheres in the form of a 3D regular polyhedron with equilateral triangles as faces (Zhi et al., 2022).

**Table 2.** Adult and Early Childhood Area Parcellations.

| Name | Number of parcels | Citation | Original Space |
| --- | --- | --- | --- |
| Gordon | 333 | Gordon, Laumann et al. 2016 | 32k_fs_LR |
| Glasser | 360 | Glasser et al. 2016 | 32k_fs_LR |
| Schaefer | 400 | Schaefer et al. 2018 | 32k_fs_LR |
| AAL | 82 | Tzourio-Mazoyer et al. 2002 | MNI152 |
| Desikan | 70 | Desikan et al. 20016 | 32k_fs_LR |
| Shen | 200 | Shen et al., 2013 | MNI152 |
| Myers-Labonte | 283 | Myers, Labonte et al., 2024 | 32k_fs_LR |
| Tu | 326 | Current study | 32k_fs_LR |
| Wang | 864 | Wang et al. 2023 | 32k_fs_LR |
| Scheinost | 87 | Scheinost et al., 2016 | MNI152 |
| Shi 1 Yr | 194 | Shi et al., 2018 | Age-specific T1<br>(Shi et al., 2011) |
| Shi 2 Yr | 205 | Shi et al., 2018 | Age-specific T1<br>(Shi et al., 2011) |
| Icosahedron<br>(control) | 304 | Zhi et al. 2022 | 32k_fs_LR |

### Comparing Our Area Parcellation to Age-specific Early Childhood Area parcellations

Using the boundary maps in Figure 2, we generated age-specific area parcellations with the BCP data divided into 5 groups and a merging threshold of 65%. To test whether finer age-specific area parcellations improve cluster validity for the corresponding age group, we calculated the DCBC for these five age-group parcellations on a secondary validation dataset containing an additional subset of BCP sessions in the same age range (N = 73 sessions from 51 participants, age range 8-29 months). This validation datset included more recently released BCP data collected at the University of Minnesota and University of North Carolina Chapel Hill sites. Acquisition and processing details were largely the same as the main BCP dataset described before with an update to the DCAN-Infant pipeline v0.0.22 where zero-padding has been implemented at the filtering step to minimize the distortions in the edges of the time series.

### Practical Implications of Using Early Childhood and Adult Parcellations

Previously, researchers have found that inaccurate area parcellations may reduce the prediction accuracy of clinical phenotypes (Abraham et al., 2017; Dadi et al., 2019). FC derived from an accurate area parcellation should yield satisfactory prediction accuracies for behavioral phenotypes (Kong et al., 2023) and demonstrate decent test-retest reliability (Tozzi et al., 2020). We thus compare the prediction accuracy of age using FC from the BCP dataset based on the present 2-year-old parcellation (Tu (326)) and the Gordon parcellation (Gordon et al., 2016), which were the best-performing early childhood and adult area parcellations on cluster validity respectively. In addition, we assessed the test-retest reliability of individual edges in the parcellated FC. We constructed a functional connectome with the first 7.2 min (600 frames for TR = 0.8 and 540 frames for TR = 0.72) of low-motion (filtered FD<0.2) fMRI data in each subject in the BCP dataset using the area parcellations and applied a linear support vector regression for the prediction of age (J. Li et al., 2024). The test-retest reliability was assessed with the first 5 min of two separate scan runs within the same session using an intraclass correlation coefficient (ICC (3,1))(Shrout and Fleiss, 1979; Tozzi et al., 2020). Details are provided in the Supplementary Materials.

### Identification of Community Structure in 2-year-olds

To characterize the relationship between the area parcels, we identified the community structure with the Infomap algorithm on the area parcels as nodes and the FC between parcels as edges (Gordon et al., 2016; Rosvall and Bergstrom, 2008). For each participant in the eLABE (Y2) dataset (N = 92), we created a parcellated time series by calculating the mean within-parcel time series over each of the parcels from the dense grayordinate time series in 32k_fs_LR space with the workbench command “wb_command cifti-parcellate”. We then cross-correlated these parcellated time series to generate a parcel-wise correlation matrix. Parcel-wise correlation matrices were Fisher z-transformed and averaged across all participants to obtain a group-average correlation matrix.

To reduce the impact of non-neuronal sources of inflation in short-distance correlation (e.g., data processing, subject motion), we applied an exclusion distance of 30 mm on the correlation matrix. A range of thresholds was then used to make the parcel-wise correlation matrix into a weighted sparse graph (edge density in steps of 0.25% ranging from 0.25% to 20%), which were entered as inputs to the Infomap algorithm. A consensus across thresholds was found with a manual examination of the communities at different thresholds to identify reliable networks across thresholds which also matched the prior description of functional systems (Power et al., 2011; Wig, 2017; Yeo et al., 2011). In addition, we also examined whether the networks at lower edge density thresholds, keeping the naming convention and colors similar to what was described in an earlier publication (Myers et al., 2024).

## Results

### Area Parcellation in 2-year-olds is Reproducible across Participants

The reproducibility of the area parcellations across participants was evaluated using split-half sampling 20 times (Figure 1A-B). We found that the reproducibility was highest around a merging threshold of 60-80%, significantly larger than the spatially permuted null model (Figure 1C-E). Based on manual inspection of the boundary map and the granularity of area parcels in popular adult area parcellations (Glasser et al., 2016; Gordon et al., 2016), we settled on a merging threshold of 65% for our main parcellation, which produced 324-391 parcels across 20 split-haves (Supplementary Figure 5A). For the remaining sections, the main area parcellation using all data in eLABE (Y2) (N = 92) and merging threshold 65% were used for evaluation, hereafter referred to as “Tu (326)”. At the merging threshold of 65%, ARI = 0.66 ± 0.02, Z-score compared to the null model = 14.3, parcel averaged dice coefficient = 0.62 ± 0.01, Z-score compared to the null model = 15.2. The dice coefficient for the binarized parcel map is 0.87 ± 0.002, Z-score compared to the null model = 8.46. Similar results were obtained with binarized boundary maps (see Supplementary Materials).

**Figure 1.**
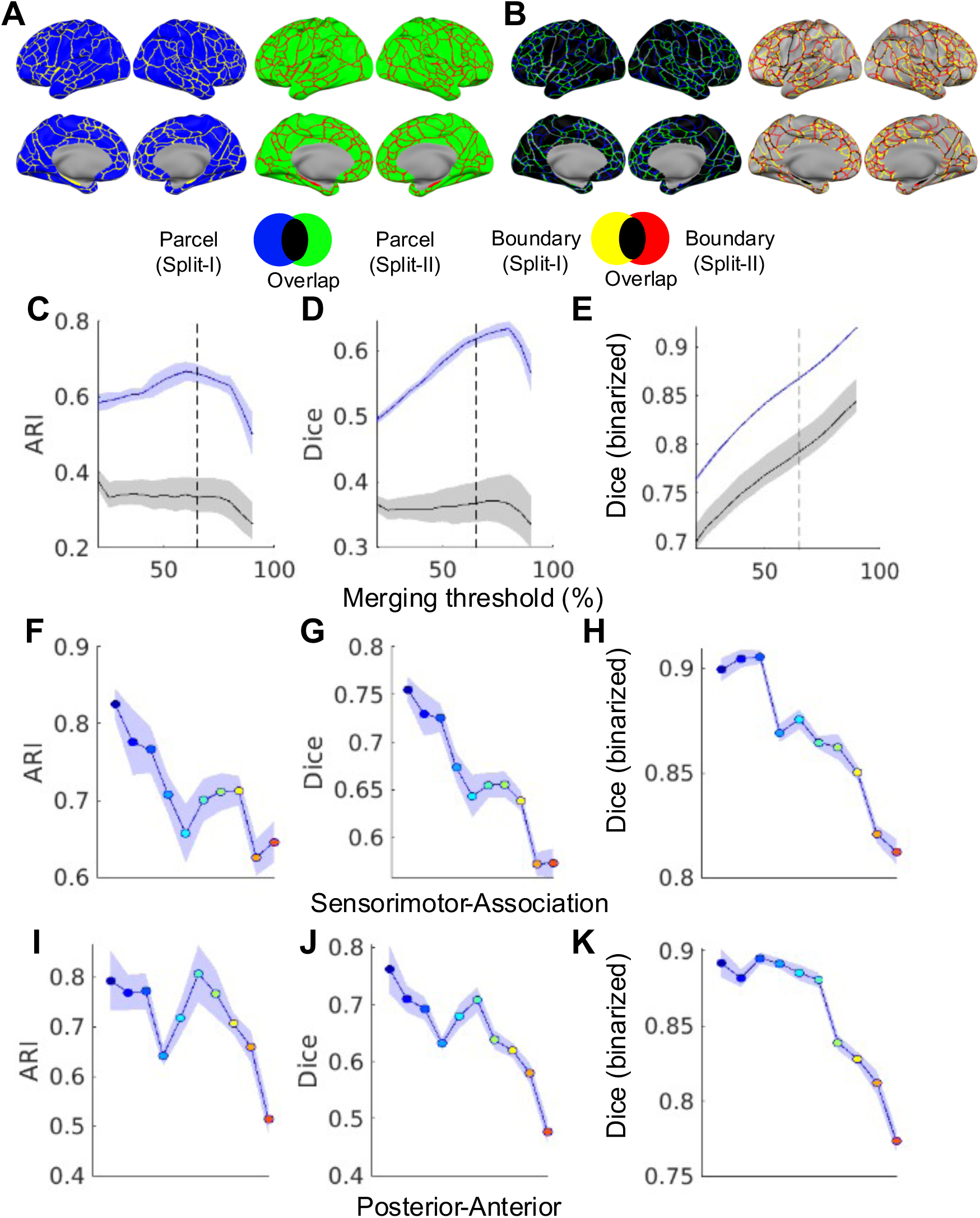
Parcel reproducibility between split halves. A) Parcellations from an example first split-half and and second split-half. B) The overlap between the parcels and boundaries in A and B. C) Adjusted Rand Index (ARI). D) parcel-average Dice coefficient. E) Dice coefficient on binarized parcels. The blue line and shaded area show the actual values and the standard deviation across 20 splits. The black line and shaded area illustrate the mean and 95% confidence interval of the spatially permuted null from one example split. The dashed line shows the merging threshold = 65%. F-H: the same metrics in C-E but separated into 10 bins along the Sensorimotor-Association axis at merging threshold = 65%. I-K: the same metrics in C-E but separated into 10 bins along the Posterior-Anterior axis at merging threshold =65%. The colors in the individual data point in F-K matches with the bin colors in Supplementary Figure 4.

Furthermore, we examined the parcel reproducibility across different positions in the brain by segmenting the brain into approximately 10 equal divisions along the sensorimotor-association axis (Supplementary Figure 4A) and the posterior-anterior axis (Supplementary Figure 4B). We found that the sensorimotor regions tend to have higher reproducibility than the association regions (Figure 1F-H) and that the posterior regions tend to have higher parcel reproducibility than the anterior regions (Figure 1I-K).

### Functional Connectivity Transition Boundaries in 2-year-olds Are Consistent Within Group and More Proximal to Those in Adults than Those in Neonates

We compared the boundary maps from the 2-year-olds (Figure 2A-B) to boundary maps generated from adults (Figure 2C) and neonates (<1 month from birth, Figure 2D) by comparing the similarity of the vertices with the top percentile of boundary probabilities (ranging from 15-55%) (Supplementary Figure 6).

**Figure 2.**
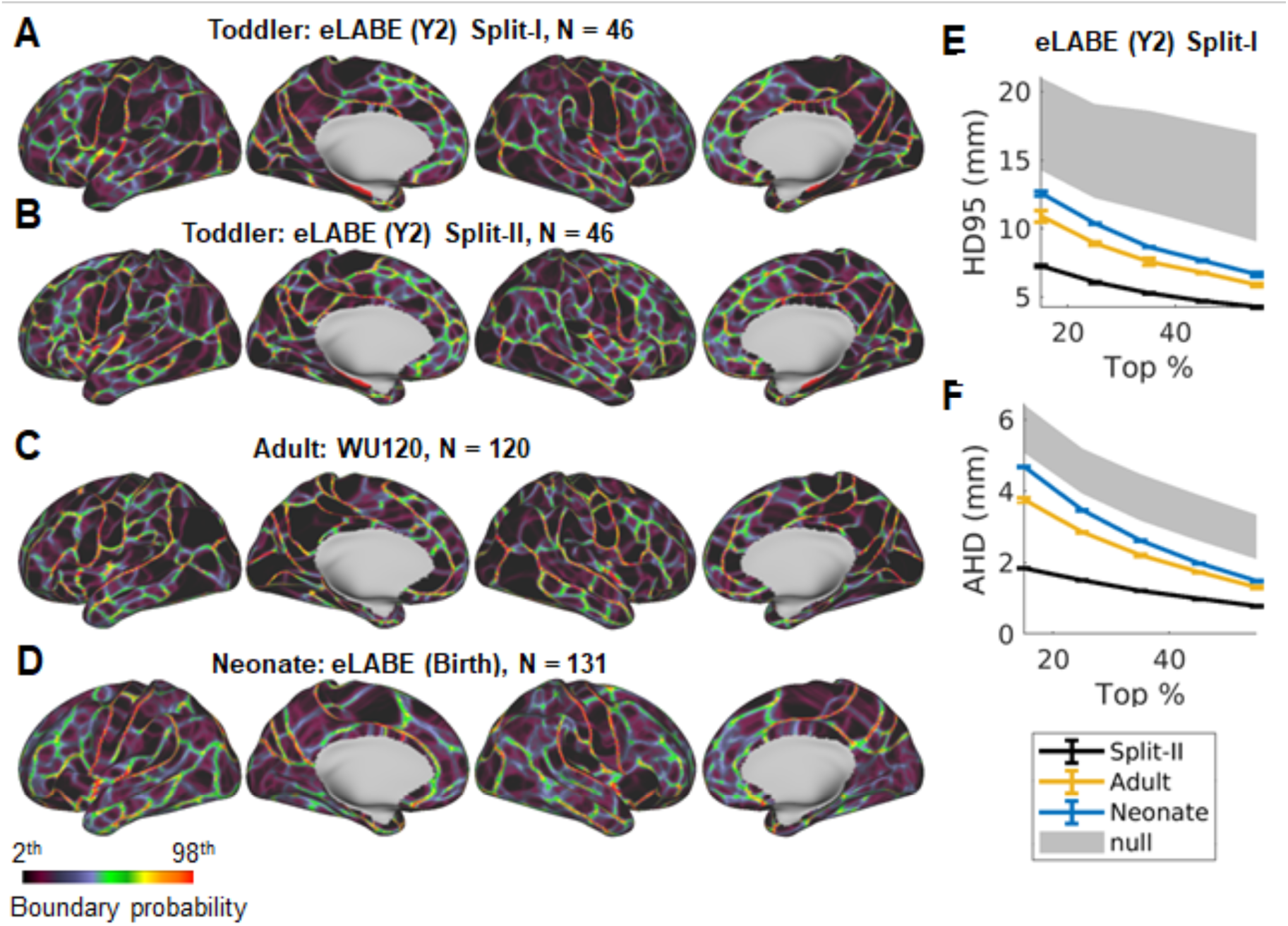
Similarity of boundary maps across ages. A) The FC boundary map in an example first split half. B) The FC boundary map in an example second split half. C) The FC boundary map in an adult dataset (WU 120). D) The FC boundary map in a neonate dataset (eLABE (Birth)). (E) 95% Hausdorff distance (HD95) indexes the spatial similarity of the boundaries between eLABE (Y2) Split-I and those from eLABE (Y2) Split-II (black), adult (yellow), and neonate (blue). The shaded area indexes the 95% confidence interval for the HD95 between the FC boundary in eLABE (Y2) Split-I and 1000 spatially permuted null of eLABE (Y2) Split-II. F) Same as E but using average Hausdorff distance (AHD). Lower HD95 and AHD indicate more similar boundaries.

Boundaries in 2-year-olds were spatially closer to adult boundaries (HD95 = 7.61 ± 0.24 mm, AHD = 2.22 ± 0.03 mm for the top 35% vertices) compared to neonate boundaries (HD95 = 8.68 ± 0.01 mm, AHD = 2.63 ± 0.01 mm for the top 35% vertices) (Figure 2E-F). The boundaries were considerably similar across the five early childhood age bins (median age 10, 12, 16, 19, 25 months, Table 1) in the BCP dataset (HD95 ≈ 5 mm for the top 35% vertices, Supplementary Figure 7). However, area boundaries tended to be more similar between early childhood groups with a smaller age difference.

### Local Gradient-Based 2-year-old Area Parcellation Provides the Best Cluster Validity for children at 1-3 years

Using FC profiles from the eLABE (Y2) individuals, we evaluated the cluster validity of the present 2-year-old parcellation versus several extant adult and early childhood area parcellations (Figure 3A), as well as a regular heaxogonal parcellation of a sphere (Icosahedron) with 304 parcels (Supplementary Figure 8) using FC from eLABE (Y2) individuals. We observed a large variation in cluster validity within adult and early childhood parcellation groups, with the Gordon parcellation demonstrating the best performance among adult parcellations and the Tu (326) parcellation demonstrating the best performance among early childhood parcellations (Figure 3B). However, all adult and early childhood parcellations examined except for AAL (82) and Desikan (70) had DCBC > 0 (FDR-corrected p<.05). The DCBC for the control Icosahedron (304) parcellation was not significantly above 0. A repeated measures ANOVA with the 13 parcellations as the within-subject factor was run on the 13×92 DCBC matrix and demonstrated a significant difference in DCBC across parcellations, F (12,1092) = 508.64, p<.001). Post-hoc paired t-test showed that Tu (326) had a better cluster validity (Cohen’s d > 2.0, Supplementary Figure 9A) than alternative adult and early childhood parcellations in eLABE (Y2) individuals.

**Figure 3.**
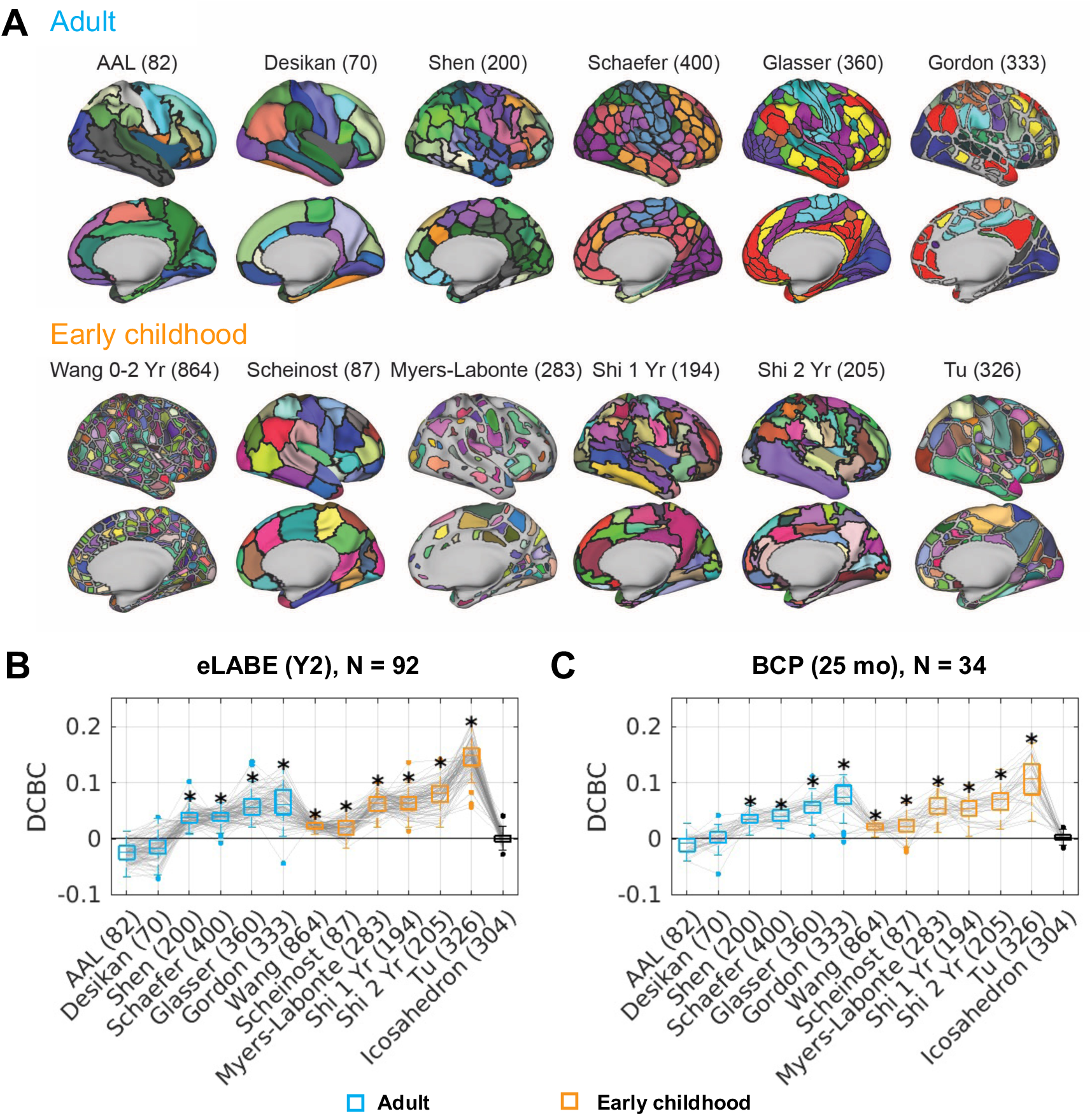
Cluster validity for different area parcellations evaluated with a distance-controlled boundary coefficient (DCBC) measure. (A) Adult area parcellations and early childhood area parcellations. (B) DCBC quantified in individuals in the same eLABE (Y2) dataset used to derive the Tu (326) parcels. (C) DCBC quantified in individuals in an independent dataset (BCP). * p<.05 after FDR correction for one-sample t-test against 0. As a convention, we noted the number of parcels of a particular parcellation scheme in parentheses, e.g., Gordon (333) means Gordon parcellation with 333 parcels

One caveat to the observation above was that the evaluation was performed on the same dataset used to generate the parcels. As such, an independent validation dataset (BCP) was used to further evaluate the cluster validity of the area parcellations (Figure 3C). The Gordon (333) and Tu (326) parcellations still performed the best within their respective parcellation age brackets, confirming the robustness of our results. A significant difference in DCBC across parcellations was found by a repeated measures ANOVA with the 13 parcellations as the within-subject factor, F (12,396) = 100.92, p<.001). Post-hoc paired t-test showed a better cluster validity of Tu (326) against other parcellations (Cohen’s d > 1.2, Supplementary Figure 9B) at 8-30 months.

To further validate the cluster validity of the parcellations in early childhood, we calculated the DCBC on individuals from all five BCP groups (Supplementary Figure 10). We ran a repeated measures ANOVA with the 12 parcellations as the within-subject factor and the 5 age bins as the between-subject factor on the 13×177 DCBC matrix. There was a significant difference in DCBC across parcellations, F (12,2064) = 551.31, p<.001), and no interaction between the five age bins and parcellations, F (48,2064) = 0.76, p = 0.88).

Similar results were observed when calculating a homogeneity Z-score at the group-average level (Supplementary Figure 11-12). Details are provided in Supplementary Materials.

### Age-specific Early Childhood Parcellations Have Comparable Cluster Validity to the 2-year-old Parcellation

We generated parcellations using the BCP dataset for five narrower age windows with a 65% merging threshold (Figure 4A). Age-specific early childhood parcellations were similar to one another (ranging from 352 to 380 parcels, ARI = 0.5-0.6). We calculated DCBC of the age-specific parcellations, the Tu (326) and the Gordon (333) on additional sessions of BCP data from a different set of subjects. A significant difference across parcellations was found with the repeated measures ANOVA with the 3 parcellations as the within-subject factor for 10 months (F(2,18) = 15.86, p<.001), 12 months (F(2,20) = 21.52, p<.001), 16 months (F(2,24) = 21.74, p<.001), 19 months (F(2,36) = 49.01, p<.001), and 25 months (F(2,38) = 48.61, p<.001). Using a post-hoc two-tailed paired t-test, we found that the age-specific parcellation outperformed Tu (326) parcellation only at 10 months (FDR-corrected p = 0.038) (Figure 4B), and was significantly worse than the Tu (326) parcellation at 19 months (FDR-corrected p = 0.0065). Given that the Tu (326) parcellation was derived from a separate dataset from the age-specific parcellations, the current results support the generalizability and the utility of our Tu (326) parcellation to the age range of 1-2 years.

**Figure 4.**
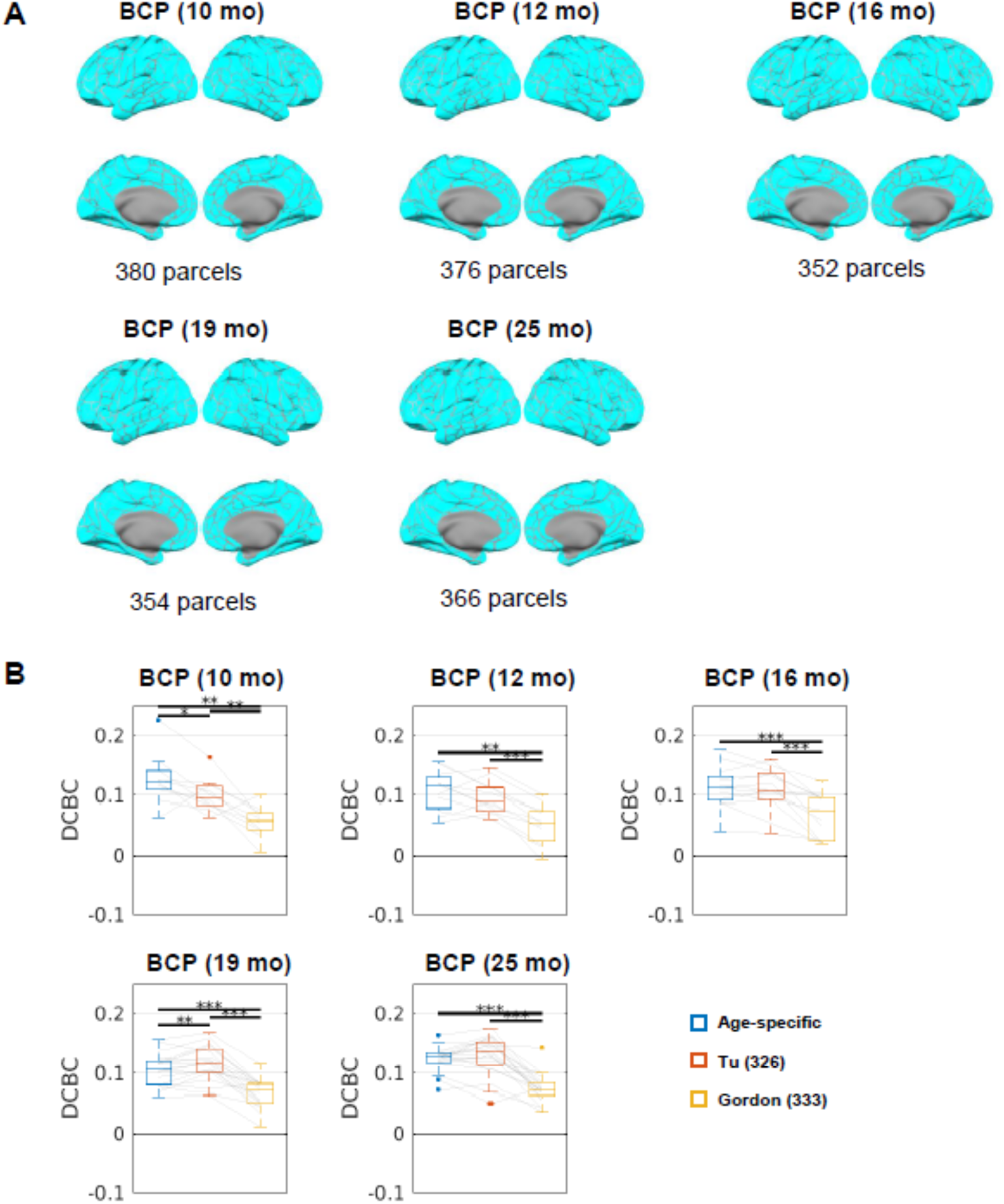
Age-specific early childhood area parcellations. DCBC on a secondary validation dataset of held-out BCP participants using i) the age-specific parcellations, ii) Tu (326), and iii) Gordon (333). ** p<.01, *** p<.001. FDR-corrected for 3 paired t-tests.

Since the Wang early childhood parcellation (Wang et al., 2023) also had age-specific versions with parcellations from 3,6,9,12,18, and 24 months, we tested whether the age-specific parcellations would best fit the individual FC in a similar age bracket. We found no clear evidence that data from a similar age range was best fit by the age-specific parcellation and that all age-specific Wang parcellations had low DCBC (<0.02) (Supplementary Figure 13).

### Adult Parcellations Based on Functional Connectivity Have a Higher Cluster Validity at Age 6 and Beyond

We determined the fit of area parcellations across the lifespan by testing our set of parcellations across FC in individual neonates (eLABE (Birth)), 3-year-olds (eLABE (Y3)), children (HBN), and young adults (HCP-YA). Neonate FC data were best fit by Myers-Labonte (283) parcellation (Figure 5A), 3-year-old FC data were best fit by the Tu (326) parcellation (Figure 5B). Children (Figure 5C-E, Supplementary Figure 14) and young adult (Figure 5F) FC data were best fit by the Gordon (333) parcellation. Adult and early childhood parcellations derived from FC rather than anatomy alone have a positive DCBC across all datasets at age 6 and beyond, with the difference in cluster validity across pairs of parcellations demonstrated in Supplementary Figure 15.

**Figure 5.**
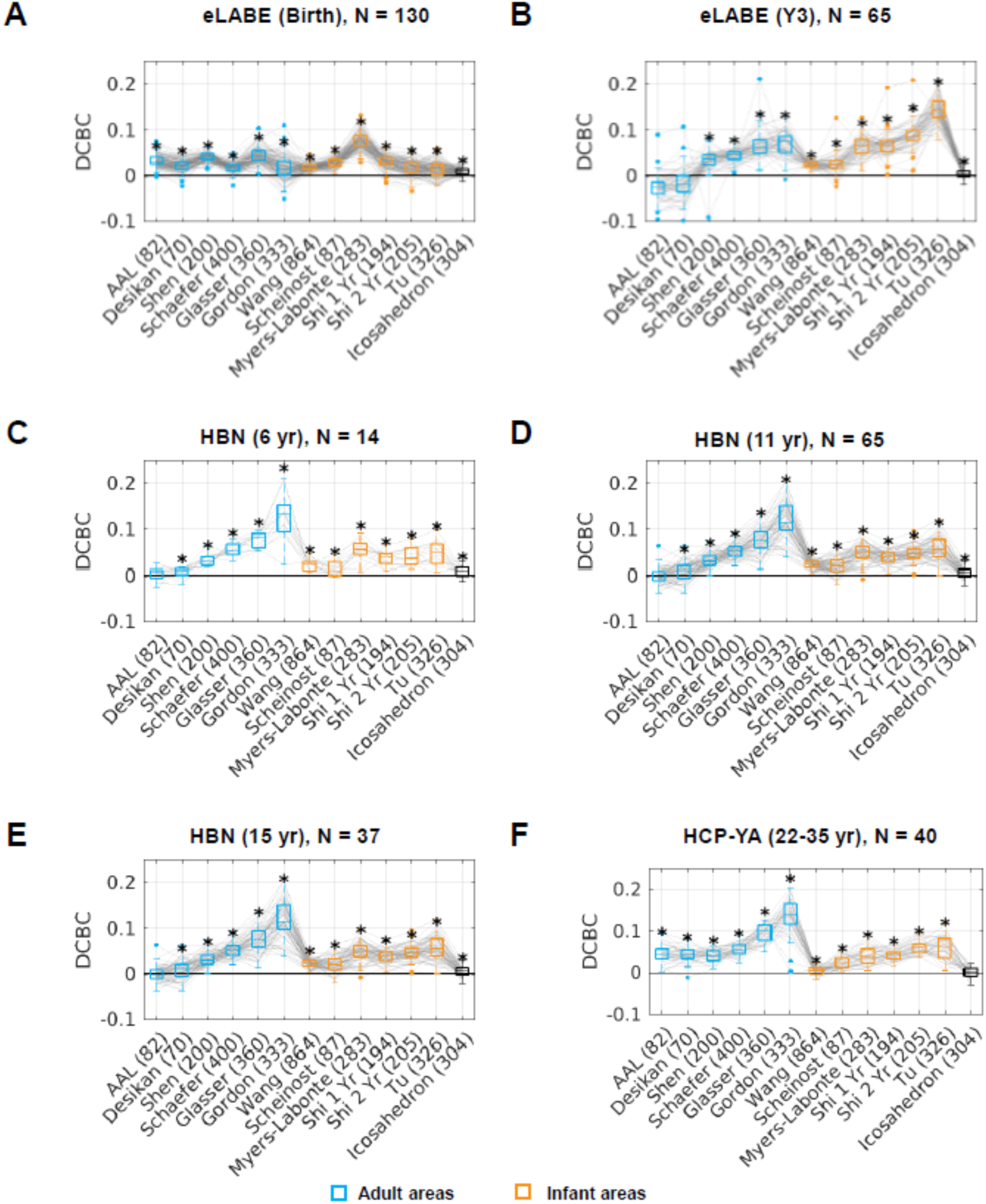
Cluster validity for different adult and early childhood parcellations across other developmental stages. A) a neonate dataset eLABE (Birth), B) an older toddler dataset eLABE (Y3), C-E) a children dataset HBN, and D) a young adult dataset HCP. * p<.05 after FDR-correction

The Myers-Labonte parcellation (Myers et al., 2024) included an alternative version that covered most of the brain (height threshold = 90%). Both versions of the Myers-Labonte parcellation significantly better fit the eLABE data at the birth time point. They were both worse than the Tu (326) parcels at the Y2/Y3 time points, and they had comparable (Myers-Labonte (283), FDR-corrected p≥.05) or worse (Myers-Labonte (370), FDR-corrected p<.05) fit than the Gordon (333) parcels at the Y2/Y3 time points (Supplementary Figure 16).

### Practical Implications of Using Early Childhood versus Adult Parcellations

To capture the practical implications of using early childhood versus adult parcellations, we tested the prediction of chronological age from the parcellated connectome and the test-retest reliability of the connectome using the validation dataset (BCP). We observed that the prediction accuracy increased with the number of parcels but plateaued at around 300 parcels (Figure 6A) regardless of adult or early childhood parcellations.

**Figure 6.**
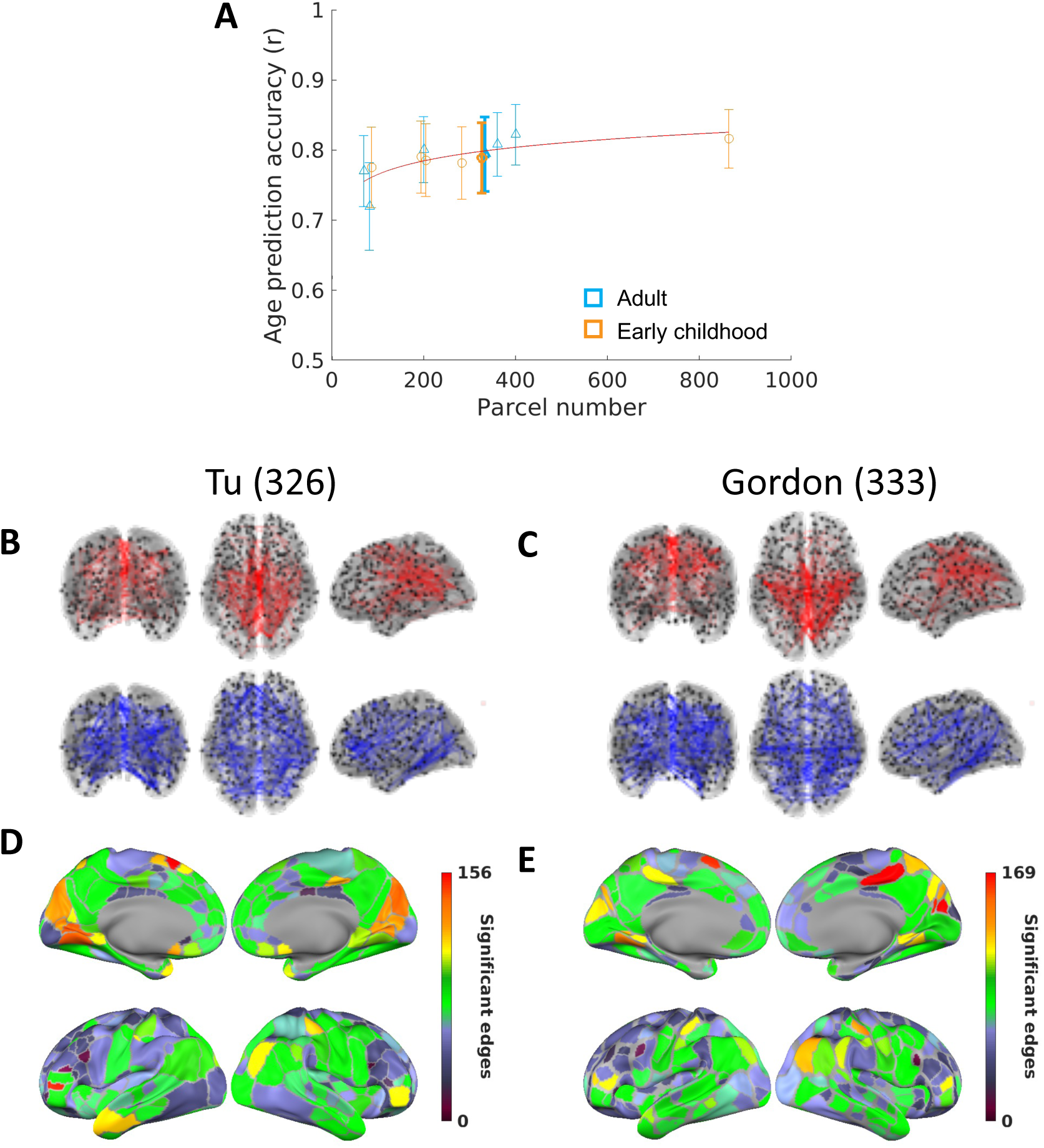
Predicting behavioral phenotypes from the connectomes using adult and early childhood parcellations. A) Multivariate prediction accuracy of age at scan using all edges. Error bars show the mean and standard deviation of correlation of actual age and predicted age at scan across 1000 random samples. The red line shows a logarithmic fit to the data. The bolded symbol shows the best adult (Gordon (333)) and best early childhood (Tu (326)) parcellations in Figure 3. B) Top 5% positive edges in age-FC correlation magnitude (red) and top 5% negative edges in age-FC correlation magnitude (blue), nodes represent parcel centroids in Tu (326) C) Same as B but for Gordon (333). D) Number of significant edges from each parcel for Tu (326). E) Number of significant edges from each parcel for Gordon (333).

The spatial distribution of the top 5% of edges with positive and negative correlations was similar across the best-performing early childhood (Tu (326)) and adult (Gordon (333)) parcellations (Figure 6B-C). Medial-visual, motor, and medial parietal areas had the highest number of edges significantly correlated with age while the lateral frontal areas had the lowest number of edges significantly correlated with age (Figure 6D-E). The spatial distribution of the top 5% of edges using other area parcellations (Figure 3) were largely consistent (Supplementary Figure 17-19), but for the coarse anatomical area parcellations, edges from some areas may appear less correlated with age (e.g. the motor cortex areas in Desikan (70)).

In addition, we computed the test-retest reliability of FC using the Tu (326) and Gordon (333) parcellations on the BCP dataset. We found lower test-retest reliability (as indexed by ICC) in the motor areas and the lateral-medial prefrontal cortex using both parcellations (Figure 7). Similar patterns were observed with other area parcellations (Supplementary Figure 20-21).

**Figure 7.**
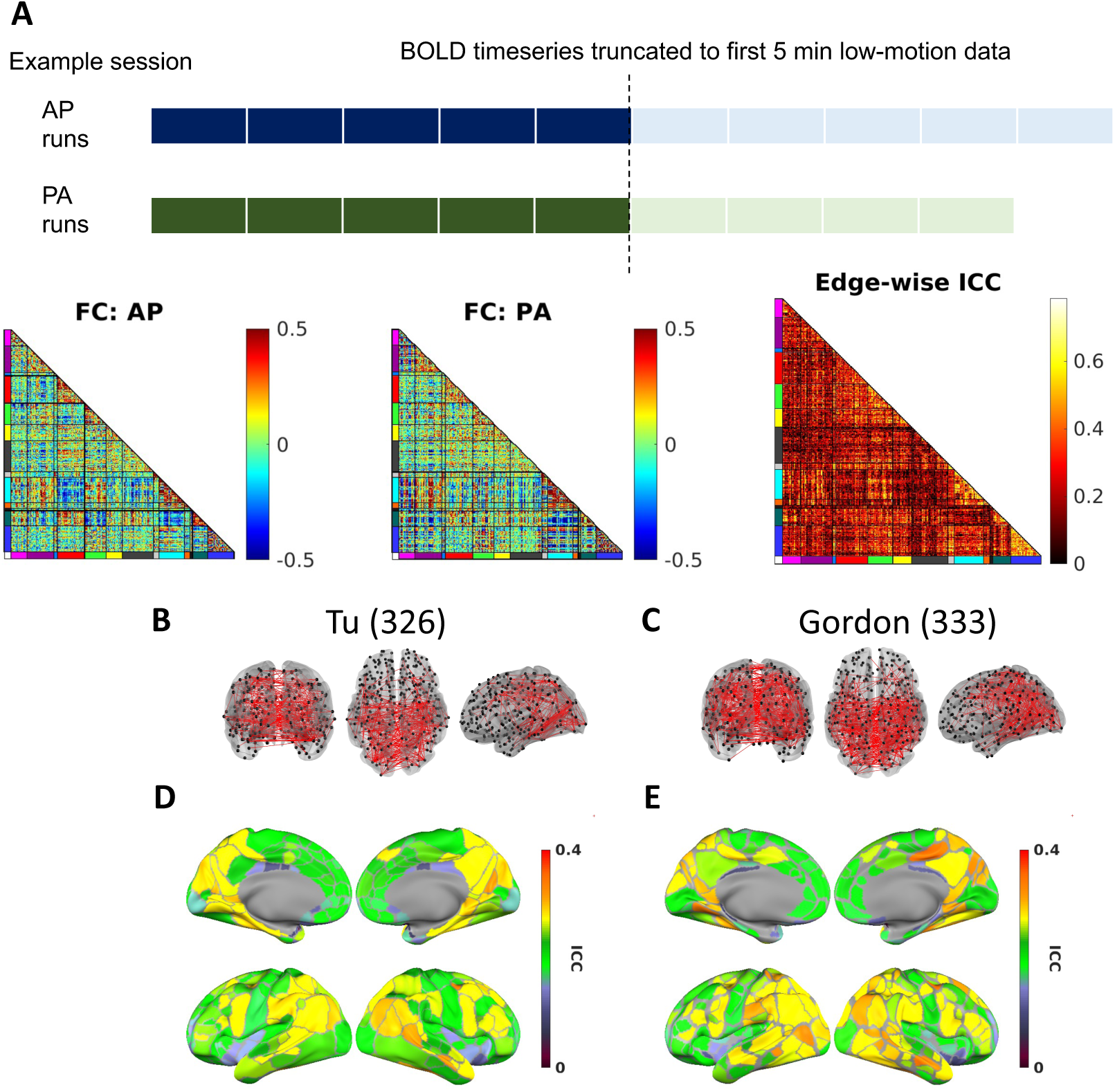
Test-retest reliability from the connectomes using adult and early childhood parcellations. A) Calculation of test-retest reliability of edges. AP = anterior-to-posterior, PA = posterior-to-anterior, example FC matrix was sorted by the Gordon et al. 2016 network orders). B) The edges with a “good” reliability (ICC = 0.60-0.75) for Tu (326). C) The edges with a “good” reliability (ICC = 0.60-0.75) for Gordon (333). D) Mean edge test-retest reliability for edges connected to areas in the brain for Tu (326). Nodes represent parcel centroids. E) Mean edge test-retest reliability for edges connected to areas in the brain for Gordon (333).

**Figure 8.**
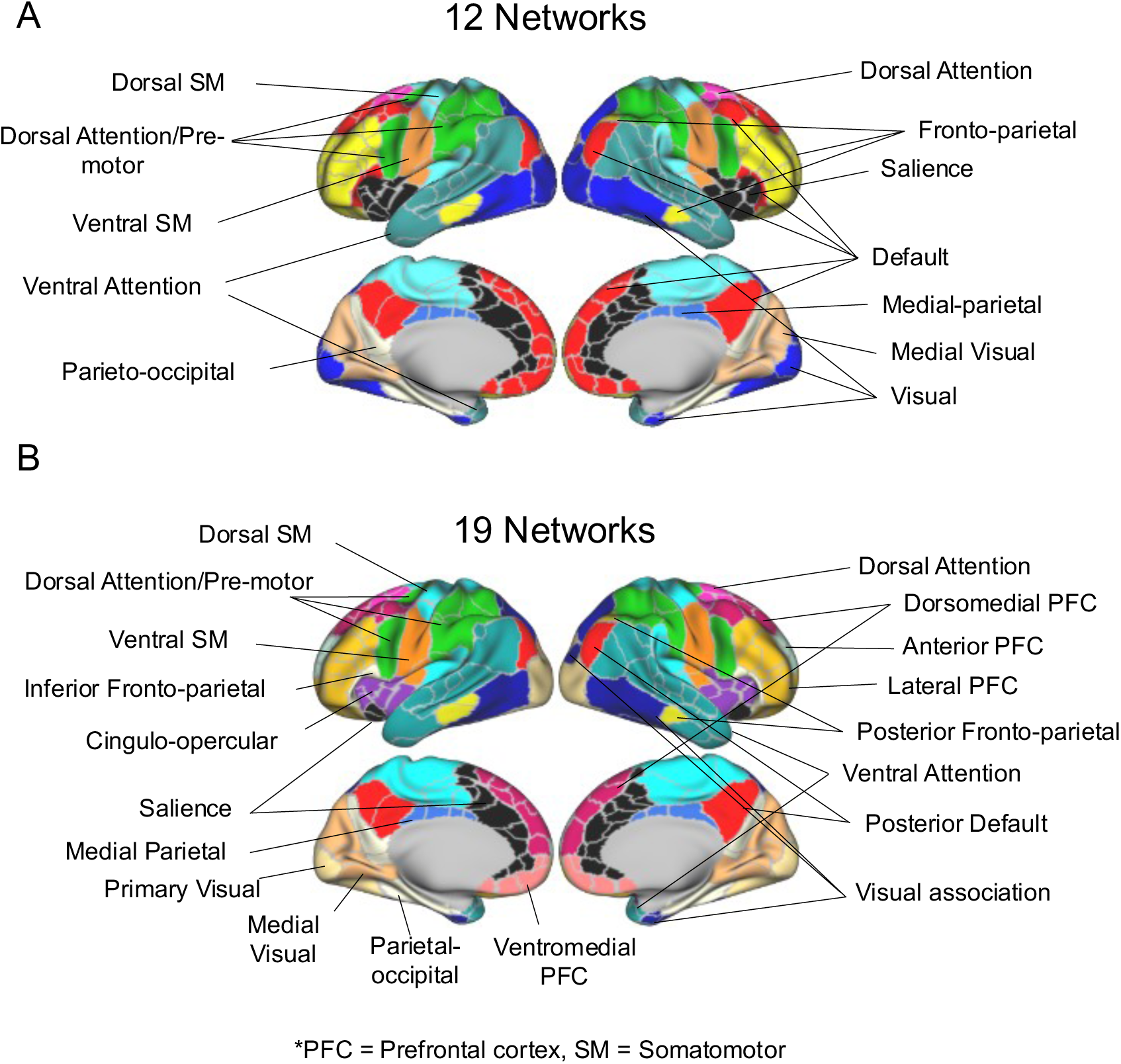
Assigned community identities for each parcel. A) Consensus community assignment for 12 networks. B) Finer division of 19 networks. Acronyms: PFC = Prefrontal Cortex, SM = Somatomotor.

### Community Assignment of Parcels into Networks

The interactions between the cortical areas form large-scale functional networks or communities (Power et al., 2011; Yeo et al., 2011). We obtained data-driven community assignments using the Tu (326) parcels as the nodes in a graph and optimized for reliable networks that were present across densities (Supplementary Video). Contrary to the fragmented anterior and posterior parts of the default network and fronto-parietal network observed in overlapping participants at birth (Myers et al., 2024; Sylvester et al., 2022), at the age of two the anterior and posterior parts of those networks joined together at higher edge densities (Figure 6A), suggestive of increased long-range FC within the network from 0 to 2 years. We found that at lower edge densities, the default network divides into four local components (posterior default, inferior fronto-parietal, dorsomedial prefrontal cortex (PFC), and ventromedial PFC) instead of distributed components (Andrews-Hanna et al., 2010; Gordon et al., 2020; Yeo et al., 2011), suggestive of more localized FC distribution in 2-year-olds compared to adults. Similarly, the fronto-parietal network can be separated into posterior fronto-parietal, lateral PFC, and anterior PFC at lower edge densities (Figure 6B). The visual network can be separated into primary visual and visual association, similar to adults. The visual association network here has sometimes been described as a component of the dorsal attention network (Du et al., 2024; Yeo et al., 2011). To illustrate the change in long-range FC strengths across neonates, 2-year-olds, and adults, we visualized the raw connectivity seed maps from different components of the canonical default network (Supplementary Figure 22).

## Discussion

We have developed a cortical area parcellation for 2-year-olds based on functional connectivity (FC). Compared to other existing adult and early childhood area parcellations, our parcellation provided a better fit for individuals between 1-3 years old across two independent datasets. Therefore, it could be used in future studies of FC in this age range. We also found that the best-performing adult parcellations also provide a good fit to 1-3 year olds, comparable to some existing early childhood area parcellations in the literature, and even outperformed our 2-year-old parcellation in children age 6 years and older.

Our parcellation achieved the best internal validity of cluster quality among all early childhood parcellations tested on the age 1-3 year-old groups. Additionally, we found that area boundaries in 2-year-olds are more similar to those in adults than to those in neonates. Taken together, these results suggest that work using adult area parcellations for developmental studies is reasonable. Our findings support the hypothesis that substantial refinement of cortical areas occurs in the first year of postnatal experience, after which they become much more adult-like. Our work not only provides insights into cortical arealization in humans during early development, but also offers practical guidelines for using cortical parcellation for neuroimaging studies involving developmental cohorts.

### Boundary Consistency in 2-year-olds Is Stronger on the Sensorimotor End Than on the Association End

We observed that area boundaries on the sensorimotor end of the sensorimotor-association hierarchy tend to be more consistent across subject samples. This observation could be attributed to two factors: 1) interindividual variability was lower in sensorimotor systems (Gratton et al., 2018; Kong et al., 2019; Li et al., 2019; Mueller et al., 2013; Sydnor et al., 2021; von Economo et al., 2008; von Economo and Koskinas, 1925), or 2) some borders in the sensorimotor systems were sharper, as seen in macaque monkeys (Lewis and Van Essen, 2000). This is consistent with the early of maturation of sensorimotor systems compared to association systems based on converging evidence including functional connectivity (Gao et al., 2015a; Smyser et al., 2011), cortical thickness (Ahmad et al., 2023), cortical surface expansion (Garcia et al., 2018; Hill et al., 2010), gray matter density (Gogtay et al., 2004), intrinsic activity (Sydnor et al., 2023, 2021), and intrinsic time scales (Truzzi and Cusack, 2023). As a consequence of an earlier maturation of sensorimotor areas, plasticity might be limited compared to association areas (Hill et al., 2010), which may lead to reduced susceptibility to environmental influences <u>(Gao *et al.*, 2017)</u> and thus lower interindividual variability in the area boundaries.

### Functional Connectivity Transition Boundaries in 2-year-olds Are More Proximal to Those in Adults than to Those in Neonates

Prior literature has described the mechanism of cortical arealization across development as a process that involved the formation of morphogen gradients driven by genetic factors, as well as the activity-dependent refinement of sharp boundaries (Cadwell et al., 2019) influenced by thalamocortical inputs (O’Leary et al., 2007). Consistent with this view, our current study demonstrates that the putative cortical areas boundaries as defined by FC transitions in 2-year-olds are more proximal to those in adults than those in neonates. These findings indicate that area boundaries experience significant development during early infancy, with the rate of change decelerating around 1 to 2 years of age.

Other studies have also identified relatively low variability in FC-gradient transition boundaries between 3 months and 24 months of age (Wang et al., 2023). This suggests that the most significant refinement of area boundaries in neonates likely occurs within the first three months. Supporting this idea, surface area continues to expand dramatically from 29 post-menstrual weeks but slows its developmental pace after three months (Bethlehem et al., 2022; Huang et al., 2022), further supporting this idea.

Notably, the area boundaries in 2-year-olds exhibit a higher similarity to those observed in neonates compared to a spatially permuted null model. This finding suggests that an organization structure of area boundaries is established from birth, aligning with the proposed intrinsic proto-mapping of cortical area organization driven by morphogens during embryonic development (Cadwell et al., 2019; O’Leary et al., 2007; Smyser et al., 2011; Tau and Peterson, 2010).

### A Coarse-grained Area Parcellation was Optimized for Biological Validity and Utility

Our level of resolution at 326 parcels is comparable to most other adult and early childhood area parcellations. In addition, it is close to the prior estimation of 300-400 cortical areas in humans based on multimodal evidences (Glasser et al., 2016; Van Essen et al., 2012; Van Essen and Glasser, 2018) for better biological interpretability. It is well-known that automated approaches are sensitive to small variations in the data, and may produce multiple solutions based on free parameter choices (Supplementary Figure 3)(Van Essen and Glasser, 2018). Since the resolution has a non-negligible effect on measurements such as global graph metrics (Arslan et al., 2018; Zalesky et al., 2010), we believe that keeping the number of parcels similar to popular adult area parcellations makes comparison across children and adults more equitable. Importantly, there are currently no 3D histological atlases showing how cortex matures during the first three years of life so a direct comparison to histological data cannot be made. While another fine-grained early childhood area parcellation exists (Wang et al., 2023), our results suggest that its generalizability to alternative processing or datasets is low as demonstrated by our results. Furthermore, multiple lines of evidence including our analyses suggest that the prediction of demographic and behavioral variables from FC data plateau with ∼300 parcels (Arslan et al., 2018; Kong et al., 2023) and that a clear correspondence between the FC gradients and the Mesulam hierarchy can be seen regardless of area parcellation scheme with more than 300 nodes (Vos de Wael et al., 2020). Therefore, having a fine-grained area parcellation may not necessarily provide a practical advantage in analyses such as examining graph properties of the brain network or multivariate age prediction.

On the other hand, we recognize that different levels of resolution may be useful in different applications (Schaefer et al., 2018; Zalesky et al., 2010). Therefore, we have also released the 2-year-old area parcellation at multiple resolutions with the caveat that our estimates of the higher-resolution area parcellations might not be as generalizable across individuals and datasets and should be used with caution.

### Cluster Validity of Adult Area Parcellations in Developmental Cohorts

We found that while the best-performing adult area parcellation (Gordon (333)) had a worse fit to the functional connectivity data in 0-3 year-olds than the best-performing early childhood area parcellations, they still had a better-than-chance cluster validity (DCBC>0) and beat the regular hexagonal (icosahedron) area parcellation. This suggests some resemblance between adult areas and areas in neonates to 3-year-olds. For additional discussion regarding results in prior literature see Supplementary Materials.

### Using Adult instead of Early Childhood Area Parcellations Lead to Qualitatively Similar Conclusions for Age Prediction and Test-retest Reliability

We found that the prediction accuracy of age increased with parcel number and plateaued around 300 parcels with no clear advantage of the shape and distribution of parcels, consistent with prior literature (Arslan et al., 2018; Kong et al., 2021). Another study found a marginal effect of atlas choice on the prediction of individual psychological and clinical traits and supported the use of data-driven rather than pre-defined area parcellations (Dadi et al., 2019).However, this observation could potentially be attributed to the difference in the number of areas between the data-driven and pre-defined area parcellations.

Consistent with prior work in adults (Tozzi et al., 2020), we found higher test-retest reliability for FC edges from the temporal and parietal lobes (Figure 7). Overall, the FC reliability in early childhood were lower than that in adults, which could potentially be explained by a combination of the low amount of data (5 min) used for test and retest, the difference in phase-encoding direction in the test and retest scans, and/or transitions between different stages of the sleep cycle in the early childhood data compared to awake adult scans (Lee et al., 2020; Mitra et al., 2017). Other studies also found relatively low edge-level test-retest reliability in infants with the mean ICC around 0.14-0.37 (Dufford et al., 2021; Wang et al., 2021).

While our limited explorations here add credence to conclusions from previous studies using adult area parcellations (Kardan et al., 2022; Nielsen et al., 2022), this does not equate to the statement that adult parcels are accurate representations of areas in early childhood.

### Network Assignments in 2-year-olds Resembled Networks in Adults

We identified fragmented components of canonical adult functional systems consistent with prior literature using similar techniques on participants within this age range (Eggebrecht et al., 2017; Kardan et al., 2022; Wang et al., 2023). This finding aligns with earlier studies suggesting that long-range FC tended to develop later than short-range FC with age (Smyser et al., 2011, 2010; Smyser and Neil, 2015; Spisák et al., 2014; Sylvester et al., 2022; Thomason et al., 2015). However, when we incorporated weaker connectivity, the network assignments in 2-year-olds had similar topography to previously reported adult networks (Gordon et al., 2016; Ji et al., 2019; Power et al., 2011; Yeo et al., 2011). Additionally, analyses from a separate study demonstrated that these network assignments provided an approximately equal fit for children aged 1 to 2 years and performed substantially better than the Gordon network assignments from young adults (Tu et al., 2024).

Despite the similarities to adult networks, we also found important differences in the network assignments in 2-year-olds. First, the temporal lobe remained largely segregated from the canonical default network unlike in adults. Additionally, the motor hand/foot system incorporated part of the inferior parietal lobule and posterior insula, which might suggest some extra plasticity that contributes to multisensory integration during development. This might be driven by the connectivity between inter-effector regions and control network (Gordon et al., 2023). Furthermore, the salience and cingulo-opercular networks were less differentiated from each other, and the cingulo-opercular network was missing the component commonly observed at the cingulate cortex.

It is important to note that the infomap community detection algorithm tends to find more localized clusters when only examining the strongest FC due to the stronger FC at short-distance, especially in developmental cohorts. We suspect that alternative methods which de-emphasize the distance dependence of FC (Sylvester et al., 2022; Zamani Esfahlani et al., 2020) may retrieve communities more similar to the large-scale functional systems identified in adults (Petersen and Sporns, 2015). Instead of making a binary decision about whether the networks “connect” or “separate”, we believe that it is more important to note the performance of the algorithm across different edge densities and compare it to the adult network topography. Thus, we provide a 12-network model which largely resembles the definition of functional networks observed in adults, and also a 19-network model with a similar granularity to the functional networks defined in neonates at birth (Myers et al., 2024) targeted at different uses.

It is worth emphasizing that whether the network clusters we identified with functional connectivity correspond to “functional systems” with specialized functional roles (Power et al., 2011; Wig, 2017; Yeo et al., 2011) remains an outstanding question. They are likely premature forms of the adult systems (Gao et al., 2015b, 2015a).The biological validity of the fragmented components we found will need to be validated with task neuroimaging data in early childhood (Yates et al., 2022, 2021) in future research. Researchers who use our network model should be fully aware of this limitation.

### Practical Recommendations on Using the Tu (326) and Alternative Area Parcellations

Age-specific parcellations are uniquely useful in that they will apply with the greatest fidelity to the age for which they were made. However, a major limitation with using an age-specific area parcellations is that a different number of nodes and edges may exist for each age. While theoretically the development of cortical areas may raise a challenge in finding a consistent area parcellation that fits all ages, our results here demonstrated that our 2-year-old area parcellation Tu (326) generalized well to fit the FC patterns in 1-to-3-year-olds. Thus, we recognize that two alternative approaches are reasonable depending on the goal and motivation of a given study.

1. *Use a canonical adult area parcellation map*. Using the same area parcellation map can ensure correspondence across age groups (Oishi et al., 2019). However, this method risks not having the best area parcellation for each group and introducing noise in the data. Based on our current results, the use of an adult area parcellation might be a reasonable choice with limited practical impact on analyses such as age prediction from parcellated connectome.
2. *Using individualized area parcellations or functional embedding to find matching relationships*. Several techniques exist to create individual parcellations based on a group-average area parcellation prior (Chong et al., 2017; Kong et al., 2021; Li et al., 2017, 2019, 2022; Qiu et al., 2022; Zhao et al., 2020), or to embed connectivity in a latent space to find correspondence across participants (Haxby et al., 2020; Langs et al., 2016; Nenning et al., 2020). Additionally, individualized area parcellations can be created with highly-sampled individuals using precision functional imaging methods (Gordon et al., 2017; Laumann et al., 2015).

### Limitations and Future Directions

Extensive work from non-human primates (NHP) has validated the neurophysiological basis of FC observed in fMRI (Logothetis et al., 2001; Pagani et al., 2023; Vincent et al., 2007), and found the fMRI-based network organization to be constrained by anatomical connectivity (Adachi et al., 2012; Pritschet et al., 2020; Vincent et al., 2007). Therefore, while functional connectivity can be affected by many factors, it indirectly provides insights into cortical arealization. However, it is important to acknowledge the inherent limitations associated with defining area boundaries based solely on fMRI (Eickhoff et al., 2015; Van Essen and Glasser, 2018). Functional connectivity estimates from fMRI do not fully account for critical neurobiological factors, including the balance of excitatory and inhibitory signals within the maturing cortex (Ben-Ari, 2002; Markram et al., 2004), the retraction of cortical fibers, and the growth or arborization of dendrites (Hua and Smith, 2004; Stiles and Jernigan, 2010). Consequently, these factors could lead to misunderstandings or misinterpretations of cortical maturation when relying solely on functional connectivity findings, and consideration of multimodal evidence could provide further insights (Glasser et al., 2016). Moreover, FC gradients identify transitions between regions that activate synchronously at rest, which correspond to features of the brain’s functional/topographical organization. For example, FC gradients segregate motor cortex into approximately foot, hand and face patches rather than elongated architectonic divisions (Glasser et al., 2016). Thus, this strategy may capture areas that are bound together by simultaneous activation (e.g., sensory and motor regions) rather than architectonic similarity (Van Essen and Glasser, 2018).

In addition, while our area parcellations described cortical area organization in 1-to-3-year-olds, future area parcellations would benefit from the additional inclusion of subcortical and cerebellar structures. Additionally, areal differentiation begins during the last part of gestation, and the current paper only considered postnatal differentiation of areas. Moreover, the group atlas can be affected by multiple factors including acquisition, resolution, consistency across participants of functional organization within areas, the consistency of system organization between areas, and the consistency of anatomic organization (Ahmad et al., 2023; Shen et al., 2013). Future research with smaller voxels or a better T2* protocol to increase signal-to-noise ratio may further improve the quality of the group area parcellation.

In addition, the datasets used for area boundaries had minor differences in acquisition and processing (Supplementary Table 1), which could potentially impact the appearance of area boundaries. In addition, future studies should also investigate how much of the differences between neonate and their older-age counterparts could be attributed to the challenges in the registration of the neonate’s brains due to their tissue properties and anatomical differences from the adult brain. Additionally, when testing the generalizability of area parcellations to neonates, 2-year-olds and 3-year-olds, we used data from overlapping subjects from the eLABE longitudinal dataset, which might have provided a slight advantage to the Myers-Labonte (283) and Tu (326) area parcellations.

One additional confound is that the early childhood data were acquired during natural sleep, which has been shown to weaken long-range connectivity within canonical functional systems (Mitra et al., 2017). Recent studies also found differences in the FC of sleeping and awake neonates (Lee et al., 2020) and infants (Tu et al., 2024; Yates et al., 2023). It is also known that the sleep architecture changes across developmental stages (Kahn et al., 1996), which may contribute to the reduced consistency of early childhood FC within and across individuals. Moreover, eyes-open versus eyes-closed seem to affect visual area functional connectivity (Laumann et al., 2015; Van Essen and Glasser, 2018). Therefore, as with all functional connectivity-based area parcellations, our putative areas may not represent exactly the neurobiological boundaries, despite its potential utility in dimensionality reduction (Van Essen and Glasser, 2018).

Lastly, some of the area parcellations tested were originally generated in volumetric space. However, all datasets used in testing the cluster validity were in the surface space. For convenience, we transformed the area parcellations in the volumetric space to a standard MNI space when necessary and then to the 32k_fs_LR surface mesh using previously described procedures (Arslan et al., 2018). This transformation was imperfect and could have unintentionally favored surface area parcellations over area parcellations which were originally determined in a volume space.

## Conclusion

We developed FC gradient-based area parcellations of the neocortical surface for 2-year-olds to be used in future studies of FC in this age range. We found that area boundaries in 2-year-olds were more similar to those in adults than those in neonates. Despite multiple similar efforts in early childhood area parcellation, our area parcellations achieved the best cluster validity among all area parcellations tested on the 1-to-3-year-olds across two independent datasets. We also found that the best-performing adult area parcellations provided a better-than-chance fit to the FC in 1-3-year-old area parcellations. Our results lend credence to conclusions from prior work using an adult area parcellation for 1-to-3-year-olds and support the hypothesis that the most substantial refinement of cortical areas occurs in the first few months of life. Our work not only sheds new insights into cortical arealization in humans but also offers practical guidelines for using area parcellations for neuroimaging studies in developmental cohorts.

## Data and Code Availability

Baby Connectome Project data are available for download at the NIH Data Repository website: https://nda.nih.gov/edit_collection.html?id=2848. Early Life Adversity, Biological Embedding (eLABE) data are available through request at https://eedp.wustl.edu/research/elabe-study/.

All analyses, unless otherwise stated, were implemented with custom MATLAB scripts in the R2020b release. All visualizations were created with <u>custom MATLAB scripts</u> or <u>Connectome Workbench</u> Version 1.5.0.

The code for the generation and evaluation of parcellations is adapted from the <u>MSCcodebase</u> and the <u>DCBC toolbox</u>. Code and parcellations mentioned in the manuscript are available to download <u>here</u>.

## Author Contributions

JCT, MDW, and ATE conceptualized the project. EMG, TOL, MM and JL provided methodology support and software. JCT and WL conducted a formal analysis. OK, LAM, EF, TKMD, AL, JKK, CS, DD, XW, and YW curated the data. JTE and CDS were responsible for project administration. MDW, JCT, JTE, DMB, BBW, JLL and CDS were responsible for funding acquisition. JCT and MDW wrote the original draft. Everyone contributed to the review and editing of the final manuscript.

## Acknowledgment

The authors thank Dr. Dustin Scheinost for providing the Scheinost parcellation in MNI space (Scheinost et al., 2016). The authors thank Drs. Zhengwang Wu, Fan Wang, and Gang Li for providing the various versions of Wang parcellation (Wang et al., 2023) in cifti 32k_fs_LRstandard mesh. The authors thank Drs. Wei Gao and Haitao Chen for their suggestions on converting the Shi parcellation (Shi et al., 2018) from infant template volume space to 32k_fs_LRsurface. The authors thank Drs. Matthew Glasser, Timothy Coalson, Caterina Gratton, Diana Hobbs, Stephanie Doerings, Gagan Wig, Da Zhi, and Richard Betzel, M. Catalina Camacho, and Scott Marek for discussions on various analyses and datasets.

During the preparation of this work the author(s) used ChatGPT in order to improve the sentence structure and language. After using this tool/service, the author(s) reviewed and edited the content as needed and take(s) full responsibility for the content of the publication.

