## Supplemental Materials for "The Generalizability of Cortical Area Parcellations Across Early Childhood"

### Supplementary Methods

#### Methods for Fast Gradient Calculations

Previous studies have demonstrated that a random subsample approximation could be used for gradient calculation to speed up the calculation and save storage with little loss in accuracy (Kong *et al.*, 2021). We therefore used a random subsampling approach for calculating and saving gradients for each participant and averaged them over participants, this allowed us to flexibly calculate the boundary maps using sub-samples of data within a manageable amount of time. This reduced computational time exponentially from creating gradients from all vertices (Gordon *et al.*, 2016). We and several researchers across institutes have validated that the resultant boundary map from the subsampling highly resembles the one from the full sample approach. Because of the stochastic nature, we also validated that the resultant maps across multiple initiations are highly similar (Pearson's r = 0.99) for the raw boundary map values. In an earlier exploration, we also discovered that this approach enables the convergence of the watershed algorithm to a similar solution across gradients with fewer steps (i.e., larger bins for each iteration of the watershed) than sampling the same vertices to calculate the gradient for each participant. This may be due to smoothing out idiosyncrasies and only retaining the most dominant and stable gradient features. This reduction in step count needed to generate consistent watershed segmentation across gradients (and hence narrow boundary maps) further reduced the computational demand. In addition, we verified that including connectivity to subcortical regions had little impact on the gradients in an adult dataset and therefore chose to omit the FC to subcortical regions for reduced computational complexity.

#### Transforming Area Parcellation to a Common 32k_fs_LR Standard Mesh

Several commonly used adult parcellations in the neuroimaging literature (Desikan *et al.*, 2006; Glasser *et al.*, 2016; Gordon *et al.*, 2016; Schaefer *et al.*, 2018) were already available in 32k_fs_LR mesh. Other cortical parcellations are only available in volume space (e.g. AAL, Shen)(Tzourio-Mazoyer *et al.*, 2002; Shen *et al.*, 2013) were previously transformed to 32k_fs_LR standard mesh (Arslan *et al.*, 2018) and obtained from GitHub (https://github.com/sarslancs/parcellation-survey-eval). According to the paper, they were "projected from volume onto the cortical surface generated from the Colin27 brain (Holmes *et al.*, 1998) using FreeSurfer (Fischl, 2012), which is then registered to the Conte69 standard space using Multi-modal Surface Matching (Robinson *et al.*, 2014). All volumetric parcellations are post-processed and each parcel is slightly dilated to fill holes that may have emerged during projection."

Similarly, several early childhood parcellations were available in 32k_fs_LR mesh with the courtesy of the authors. Other early childhood area parcellations originally only available in volumetric space were manually transformed in-house as described below: The Shi and Scheinost parcellations (Scheinost *et al.*, 2016; Shi *et al.*, 2018) were provided in the MNI volume space by the authors and were manually transformed to surface space using the Conte69 midthickness 32k_fs_LR surfaces using the

Connectome Workbench command: “wb_command –volume-label-to-surface-mapping”. The resulting file was dilated using “wb_command –cifti-dilate” with 10mm to remove all holes from the volume-to-surface transformation. Next, the isolated patches with a connected component size smaller than 0.1 of the average parcel size of each hemisphere are replaced with the mode of their neighboring vertices using the code from the CBIG repository (https://github.com/ThomasYeoLab/CBIG/tree/master/utilities/matlab/parcellation/CBIG_CleanSurfaceParcellation.m). A final step removes parcels smaller than 10 vertices. We tried to retain the faithful mapping of the original volumetric parcellation as best as we could, but this procedure does have a potential limitation of reducing the performance of the parcellation on evaluation metrics.

**Measures of Cluster Validity**

Global homogeneity of FC (Craddock *et al.*, 2012; Gordon *et al.*, 2016) was one common measure used to quantify the cluster validity. The homogeneity of the parcels was calculated as the variance explained by the first principal component of the FC profiles across all vertices in a parcel (Gordon *et al.*, 2016). Homogeneity of the parcellation was sometimes calculated by the average homogeneity across all parcels (Gordon *et al.*, 2016; Myers *et al.*, 2024). However, this metric tended to be biased towards parcellations with a skewed parcel size distribution (Arslan *et al.*, 2018), so we also calculated the parcel-size weighted average homogeneity (Arslan *et al.*, 2018; Schaefer *et al.*, 2018), where the weight was determined by the ratio of vertices in each parcel and the total vertices in all valid parcels. To obtain a null model for global homogeneity, we inflated the cortical surfaces to spheres and applied a random rotation around the x, y, and z axes 1000 times, before mapping the rotated coordinate to the cortical surfaces (Gordon *et al.*, 2016). This operation often creates missing data by generating rotations into the medial wall and may also change the size of the parcels slightly hence creating a bias (Markello and Misic, 2021). Additional steps to deal with the missing data such as imputation, and dilating or contracting the rotated parcels to match the actual size of the parcel were implemented, and low SNR region was also excluded from the homogeneity calculation (Gordon *et al.*, 2016; Myers *et al.*, 2024). In contrast, prior studies (Myers *et al.*, 2024) removed the parcels from the homogeneity calculation if they had <15 vertices outside the low SNR regions, rotated the remaining parcels, removed the rotated parcels if they ended up in the medial wall or if the parcels had ≥ 15 vertices inside the low SNR regions. The homogeneity of the removed parcel during rotations was imputed by the mean value across other rotations. We used a dataset-specific low SNR map determined as the average of session-mean BOLD signal across participants <750 in BOLD 1000 normalized data for the eLABE (Y2) data as our low SNR regions. We calculated the homogeneity following three different procedures: A) the same as reported in prior literature (Myers *et al.*, 2024), B) excluding the actual and rotated parcels if they had ≥ 15 vertices inside the low SNR regions, C) not considering the low SNR regions.

Recent studies showed that global homogeneity did not account for the intrinsic spatial smoothness of the data, and thus random but spatially contiguous parcellations tended to achieve high global homogeneity (Zhi *et al.*, 2022). Even though in theory this could be controlled for with a spatial permutation test by calculating a Z-score compared to a rotated null model, it was subject to the limitations mentioned above. In addition, prior studies only tested the cluster validity using homogeneity Z-score on group-average functional connectivity data, rather than assessing the fit to individual data, and assessing the effect size and statistical significance of the cluster validity across parcellations in a given cohort. Here, we adopted a distance-controlled boundary coefficient (DCBC) measure which attempted to minimize the spatial bias directly by binning the FC profiles by distance and calculated the similarity of the FC profiles within and between parcels for each given distance bin (e.g., between 10mm and 11mm) (Zhi *et al.*, 2022).

We calculated DCBC with our custom MATLAB scripts closely following previously published methods and code (https://github.com/DiedrichsenLab/DCBC) using the default parameters: 1mm in size and 35 bins for a balance between bias, accuracy, and computational efficiency. We used the geodesic distance of the Conte69 atlas to calculate the distance bins. On top of this, there are two minor modifications that we considered to be more appropriate for the metric. First, we Fisher-Z-transformed the correlation values before averaging them and applied an inverse Fisher-Z-transformation at the end instead of directly averaging the correlation values in each bin. Secondly, we excluded the vertices with no BOLD signal in the session when considering between-parcel profiles.

**Prediction of Behavioral Phenotypes**

Age and other developmental scores can be predicted from an individual's functional connectome using simple machine-learning models. We explored whether the area parcellation choice affects the accuracy of this prediction in the BCP dataset. Specifically, we employed a multivariate prediction using a linear support vector regression (LSVR) model (Smola and Schölkopf, 2004; Cui and Gong, 2018) with ridge regularization and split the subjects into a training set and test set 1000 times. For each round of 1000 random samplings, 80% of the subjects were entered into the training set, and 20% of the subjects were entered into the test set. To avoid potential problems caused by shared variance, correlated samples (i.e., multiple sessions of the same subject) were always kept in either the training or test set. For age prediction, we have 177 sessions from 112 unique subjects. The regularization parameter lambda in LSVR model was tuned by an embedded 5-fold cross-validation (CV) in the training set from $10^{-1}$ to $10^{-5}$ equally spaced in log space by minimizing the mean square error (MSE). The trained LSVR model with the optimized hyperparameter was used to predict the age from the test set using the functional connectomes from the test set. The Pearson correlation (r) between the predicted and actual ages along with the MSE was calculated as the measure of prediction accuracy.

**Test-retest Reliability**

As described in prior literature, we quantified the test-retest reliability of FC with intraclass correlation coefficient (ICC) as implemented in MATLAB (https://www.mathworks.com/matlabcentral/fileexchange/22099-intraclass-correlation-coefficient-icc). In particular, we assessed the consistency among measurements under the fixed levels of the session factor (Tozzi *et al.*, 2020). This measure has been named ICC 'C-1' (McGraw and Wong, 1996) or ICC (3,1) (Shrout and Fleiss, 1979).

For this analysis, we used the BCP dataset for the 66 sessions with at least 5 minutes of low-motion data (FD<0.2) for both AP and PA acquisition directions. We defined the first 5 min of low-motion data from scans in AP direction as "test" and PA direction as "re-test". Two FC matrices were constructed, one from the AP runs and one from the PA runs (Supplementary Figure 18A).

To calculate ICC for all our FC values, first, the FC values in the upper triangle of each subject's connectivity matrix were entered as rows in two large matrices (one matrix for "test" and another for "re-test", one row per subject in each matrix). Then, the corresponding columns of these matrices were compared to obtain an ICC value for each edge.

### Supplementary Results

#### Reproducibility of Binarized Boundary Maps

We found the binarized boundary maps (parcels as 0 and boundaries as 1)(Han et al., 2018; Wang et al., 2023) were also more similar across the split halves than the spatially permuted null model, with a higher dice coefficient (0.35 $\pm$ 0.007, Z-score = 11.8), a lower HD95 (6.90 $\pm$ 0.32 mm, Z-score = -5.5), and a lower AHD (2.07 $\pm$ 0.07 mm, Z-score = -7.06) at merging threshold = 65% (Supplementary Figure 5B-E).

### Cluster Validity Results with Homogeneity and Its Relationship to DCBC

Our preliminary explorations suggest that choices to deal with low SNR regions in the homogeneity analysis using a spatial permutation null can impact the conclusions drawn about parcellation performances. For example, having a more stringent criterion for removing rotated parcels compared to actual parcels as described above tends to favor the null hypothesis. Additionally, when the global homogeneity of a parcellation was calculated with the unweighted arthmetic average of the homogeneity in individual parcels, the parcellation homogeneity would be higher for a parcellation with many small parcels and a few large parcels compared to one with similarly sized parcels.

In general, the DCBC metric and the homogeneity Z-score calculated with three different procedures to deal with low SNR regions demonstrated a positive correlation (Supplementary Figure 12-13). With all three methods, the DCBC is correlated with the homogeneity Z-score, especially when homogeneity is calculated as a parcel-size weighted average across all parcels (Supplementary Figure 13).

In conclusion, while taken at face value, our results in Figure 5A contradicts with an earlier study which suggested that most adult parcellation had worse than chance homogeneity fit (Myers *et al.*, 2024), this might be due to the more conservative choice of null models in the prior study than the current study, a.k.a. having a more stringent exclusion criterion for rotated parcels compared to actual parcels. In addition, our findings compared the parcellation fit in individuals rather than a group-average. Therefore, while our results generally agree with the prior conclusions, the assertion that adult parcellation do not fit the neonate data at all might be a little too conservative.

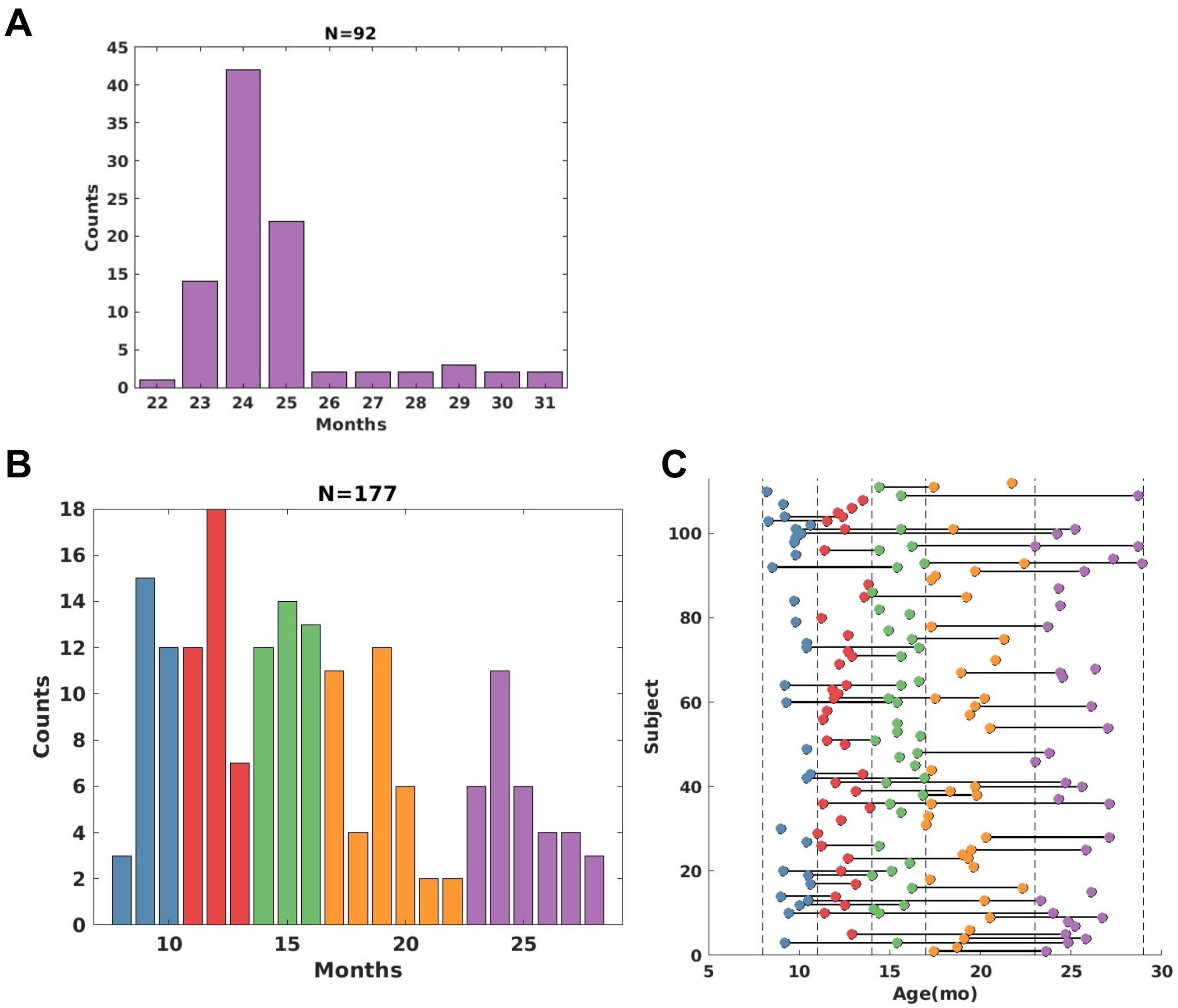


**Supplementary Figure 1.** *Subject demographics for eLABE (Y2) and BCP datasets.* A) The 92 sessions from independent subjects in eLABE (Y2). B) The 177 sessions from 112 subjects in BCP were divided in 5 groups of 30-40 sessions per group. C) The longitudinal sessions for each of the 112 subjects in BCP.

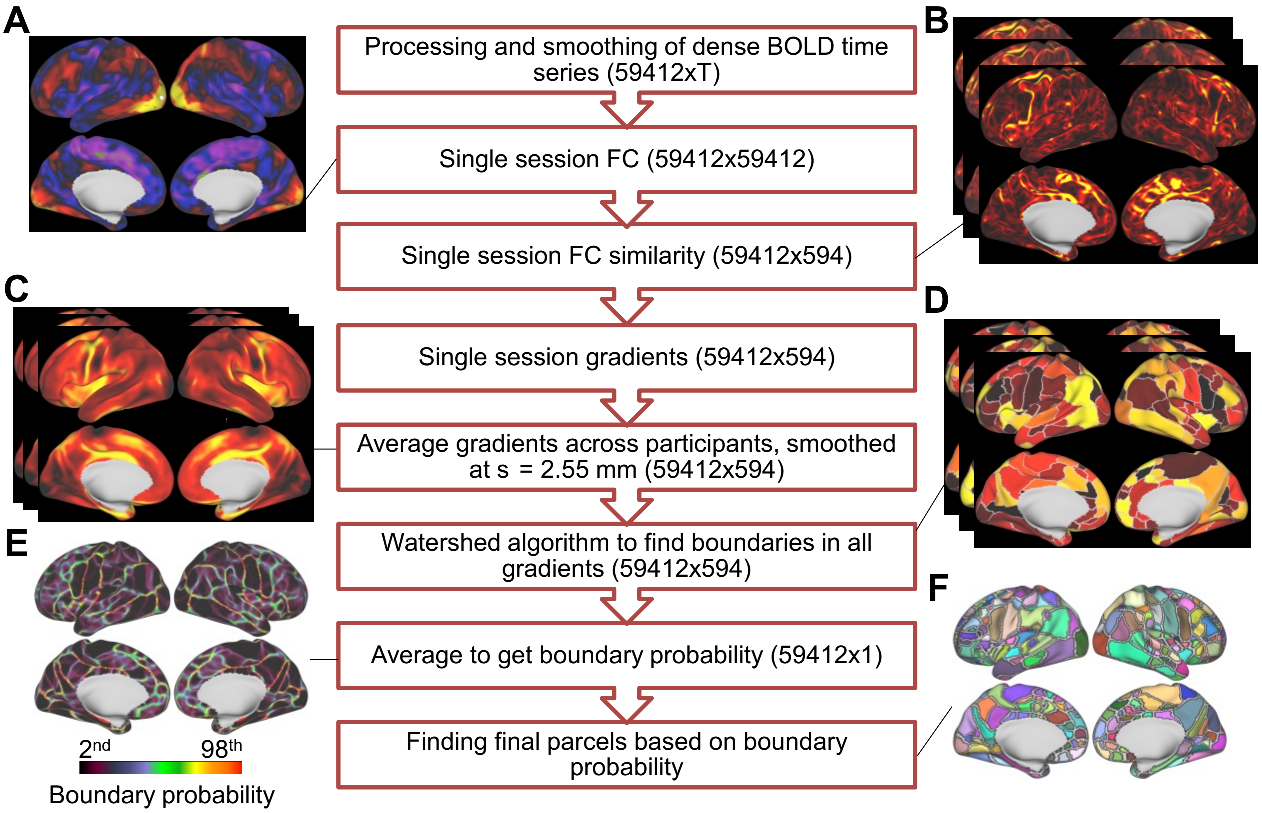


**Supplementary Figure 2.** *Creating toddler area parcellation with resting-state functional connectivity boundaries using 92 participants at age 2 (eLABE (Y2))*. A) Example FC with a seed in the visual cortex. B) Example FC similarity based on the similarity of FC at each vertex to that of 594 randomly placed seeds. C) Averaged gradient across participants. D) Putative boundaries detected from each gradient using the watershed image segmentation algorithm. E) Boundary probability maps calculated by averaging the boundaries in D. Green/red color represents a high probability of boundaries while black color represents a low probability of boundaries. F) Discrete parcels from the boundary probability in E using a merging threshold of 65%.

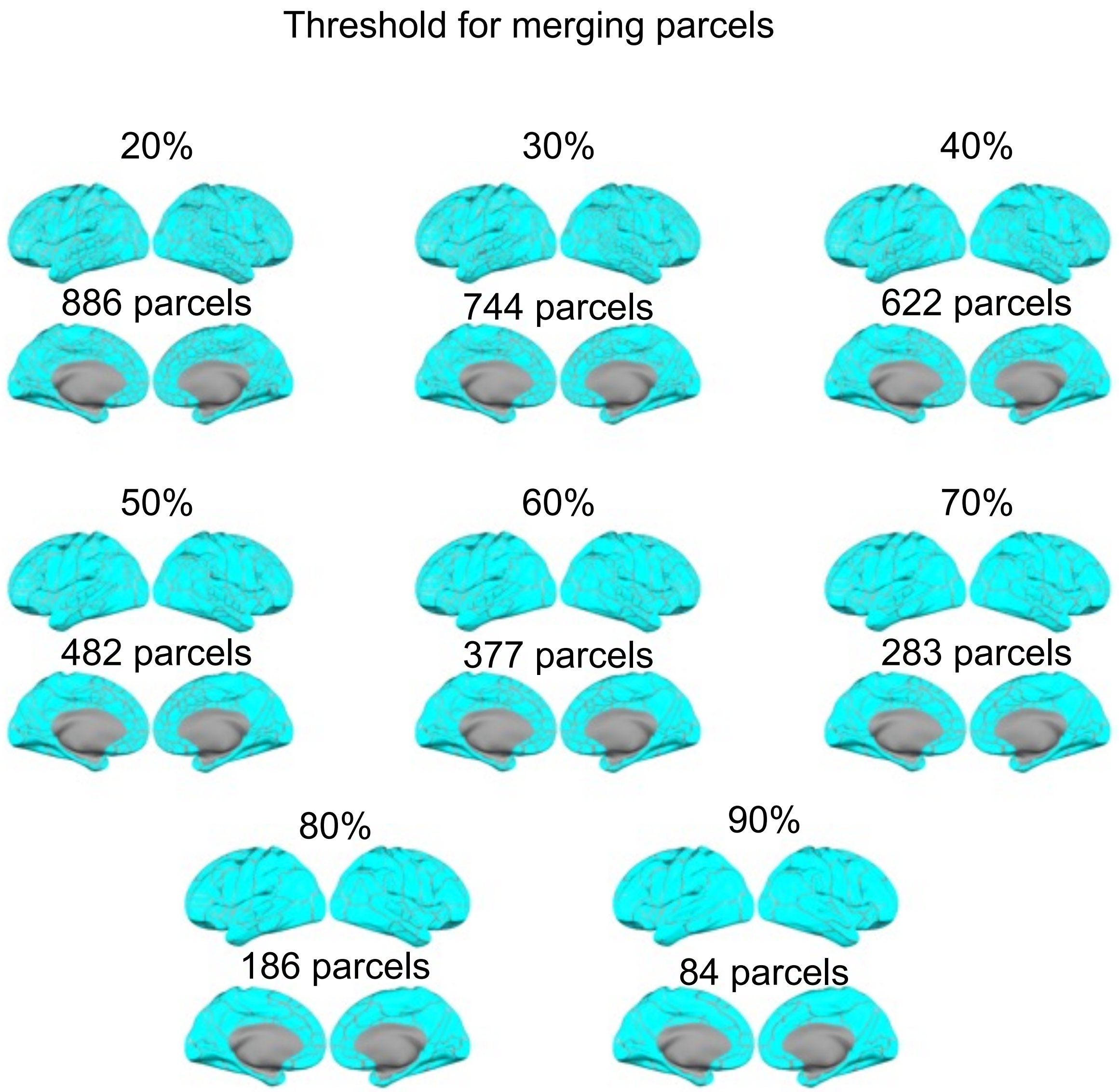


**Supplementary Figure 3.** *Toddler parcellations at different merging thresholds.*

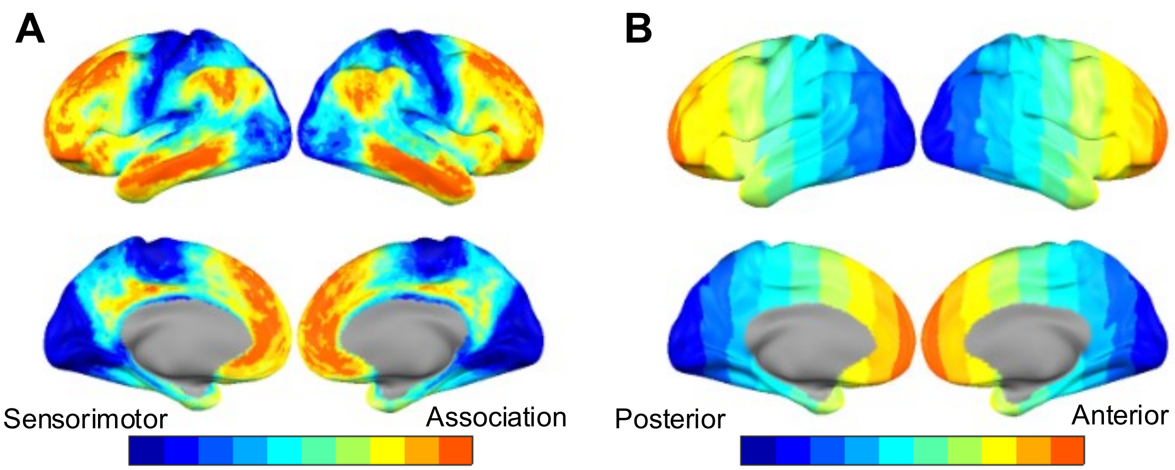


**Supplementary Figure 4.** *Division of the cerebral cortex into 10 bins by* A) the Sensorimotor-Association axis, B) the Posterior-Anterior axis.

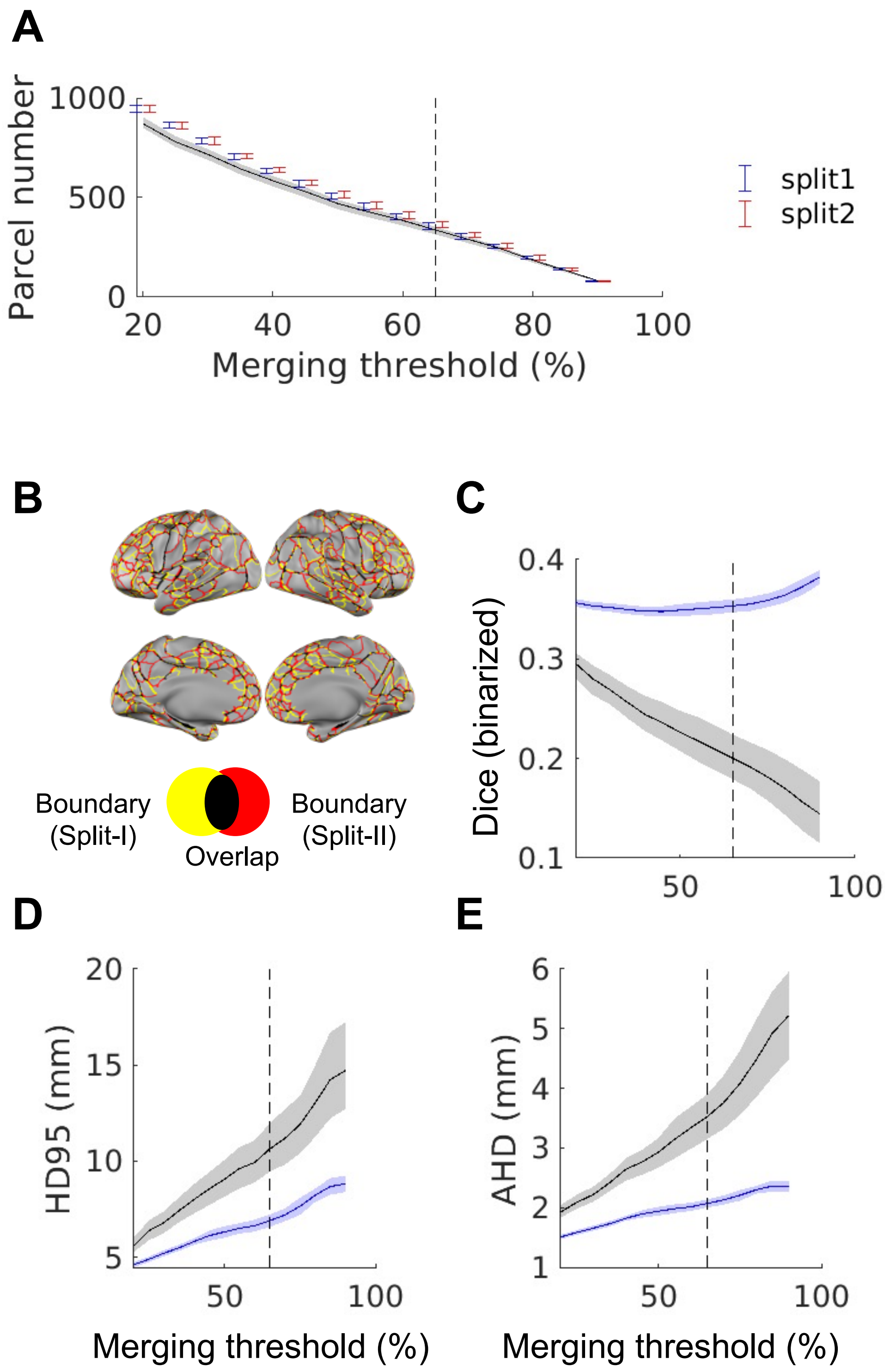


**Supplementary Figure 5.** *Parcel size and boundary similarity for different merging thresholds.* A) Parcel number across merging threshold. The error bars demonstrate the mean and standard deviation across 20 splits. The blue line and shaded area show the actual values and the standard deviation across 20 splits. B) Parcel boundaries from two non-overlapping split haves. C) Dice coefficient for binarized boundaries. D) 95% Hausdorff distance (HD95). E) average Hausdorff distance (AHD). The black line and shaded area illustrate the mean and 95% confidence interval of the spatially permuted null from one example split. The dashed line shows the merging threshold = 65%.

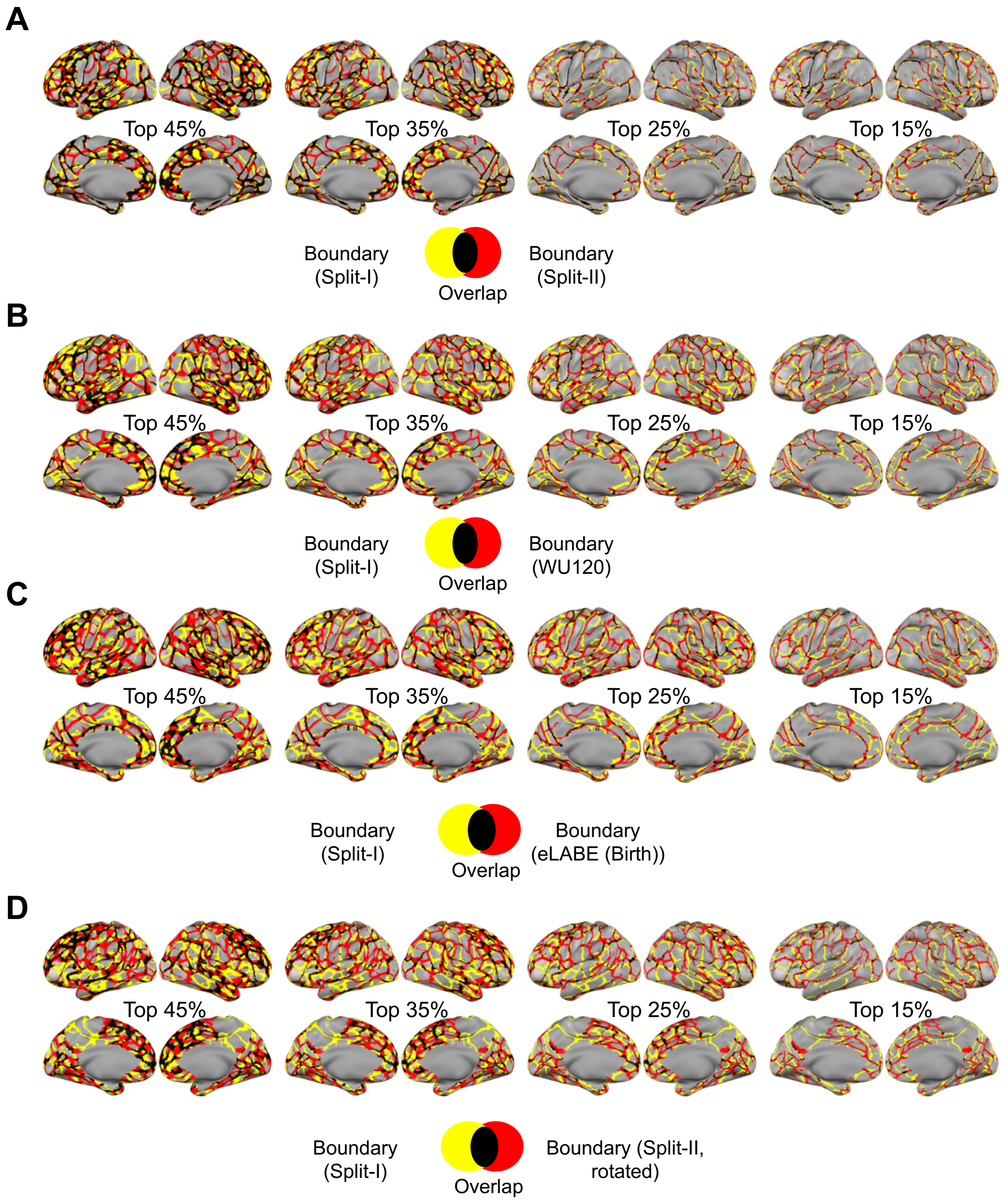


**Supplementary Figure 6.** *Overlap between the top boundary probability*. A) Comparing with the top boundaries from Split-II. B) Comparing with the top boundaries from WU120. C) Comparing with the top boundaries from eLABE (Birth). D) Comparing from the top boundaries from Split-II, rotated.

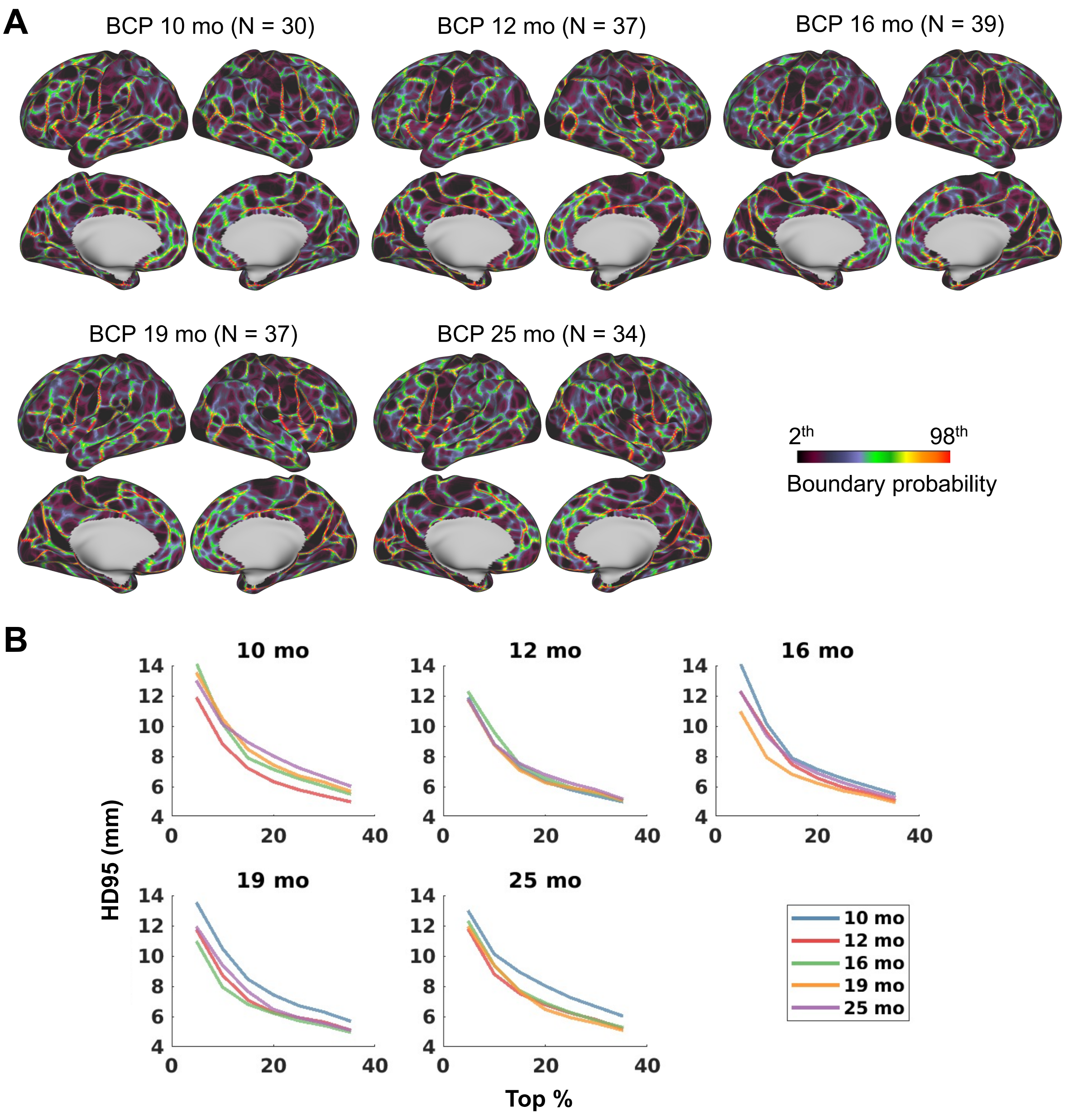


**Supplementary Figure 7. Similarity of the boundary maps across infancy to toddlerhood in the validation dataset (BCP).** A) The boundary probability maps for the 5 groups, B) The 95% Hausdorff distance (HD95) in mm between the boundaries with 5-35% of the top vertices.

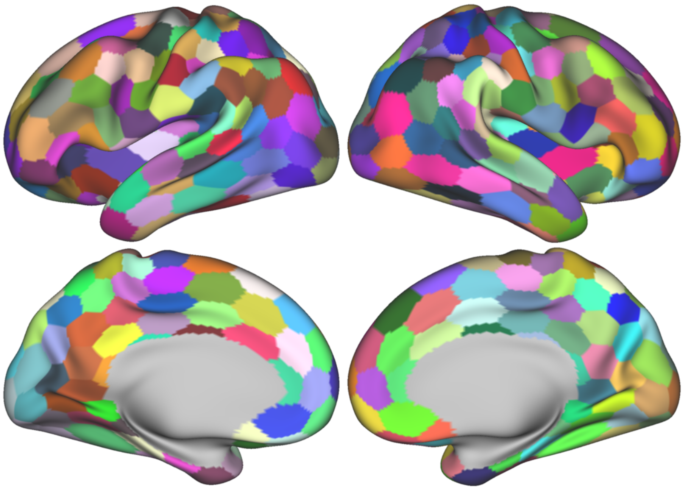

**Supplementary Figure 8.** Icosahedron parcellation with 304 parcels.

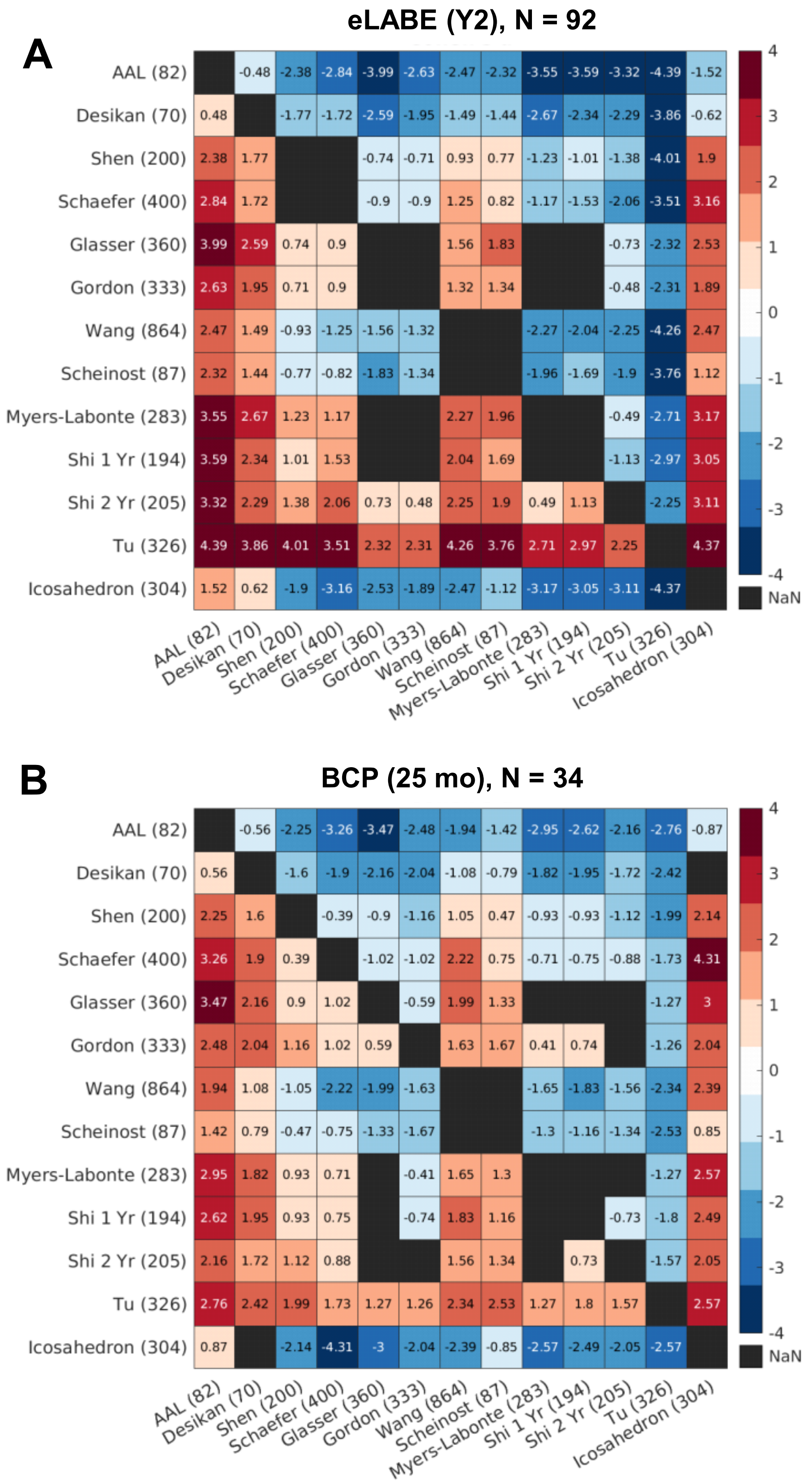


**Supplementary Figure 9.** *Effect size (Cohen's d) of the difference between parcellations in 2-year-olds datasets.* A) eLABE (Y2). B) BCP (25 mo). Each block shows the comparison between the parcellation in each row against the parcellation in each column. Non-significant differences were displayed as black blocks (NaN).

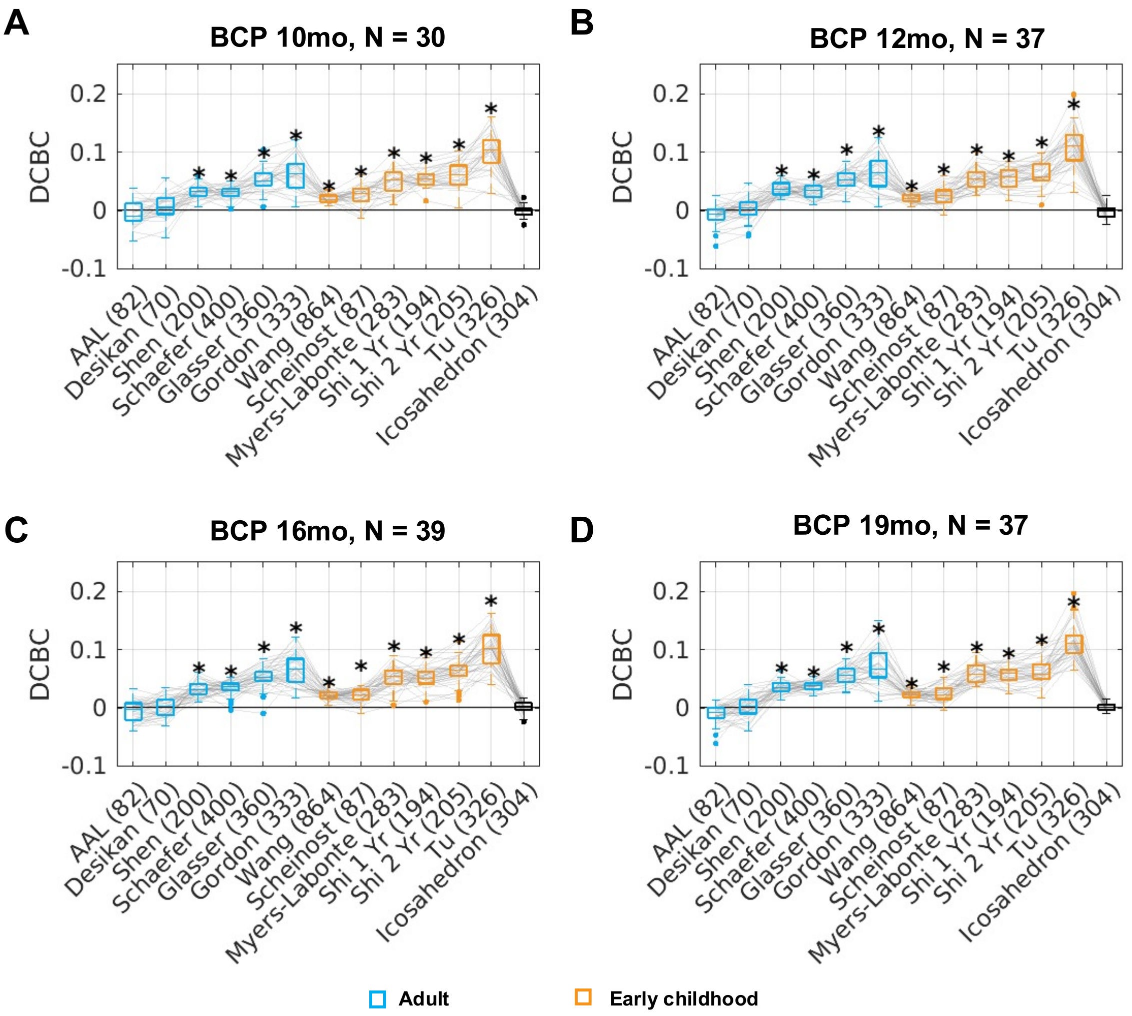


**Supplementary Figure 10.** *Distance-controlled boundary coefficient (DCBC) cluster validity for different adult and early childhood area parcellations for other groups than 25 mo in the BCP dataset.*

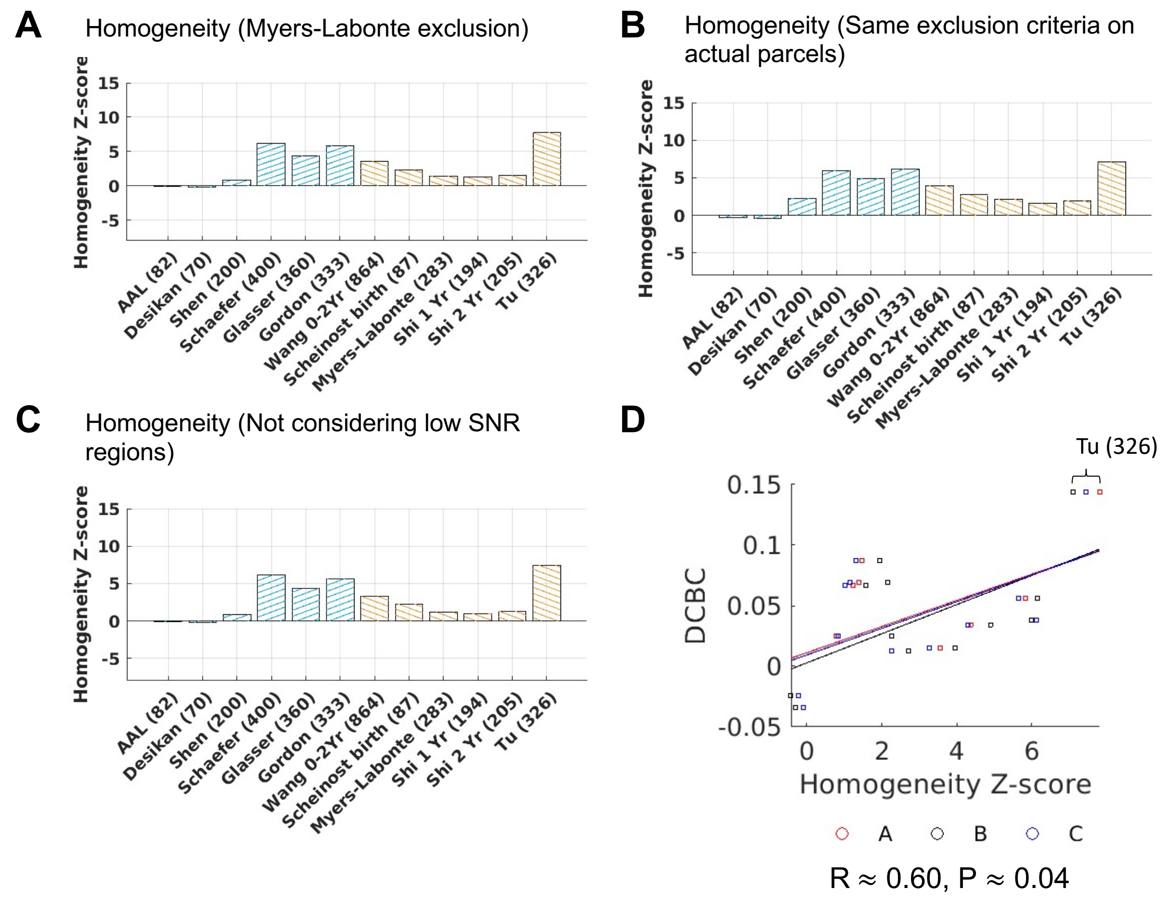


**Supplementary Figure 11.** *Functional connectivity homogeneity Z-scores using adult and early childhood parcellations on group-average eLABE (Y2) data (average homogeneity calculated without weights).* A) Homogeneity Z-score (Myers-Labonte exclusion). B) Homogeneity Z-score (Same exclusion criteria on actual parcels). C) Homogeneity Z-score (Not considering low SNR regions). D) DCBC against homogeneity Z-score.

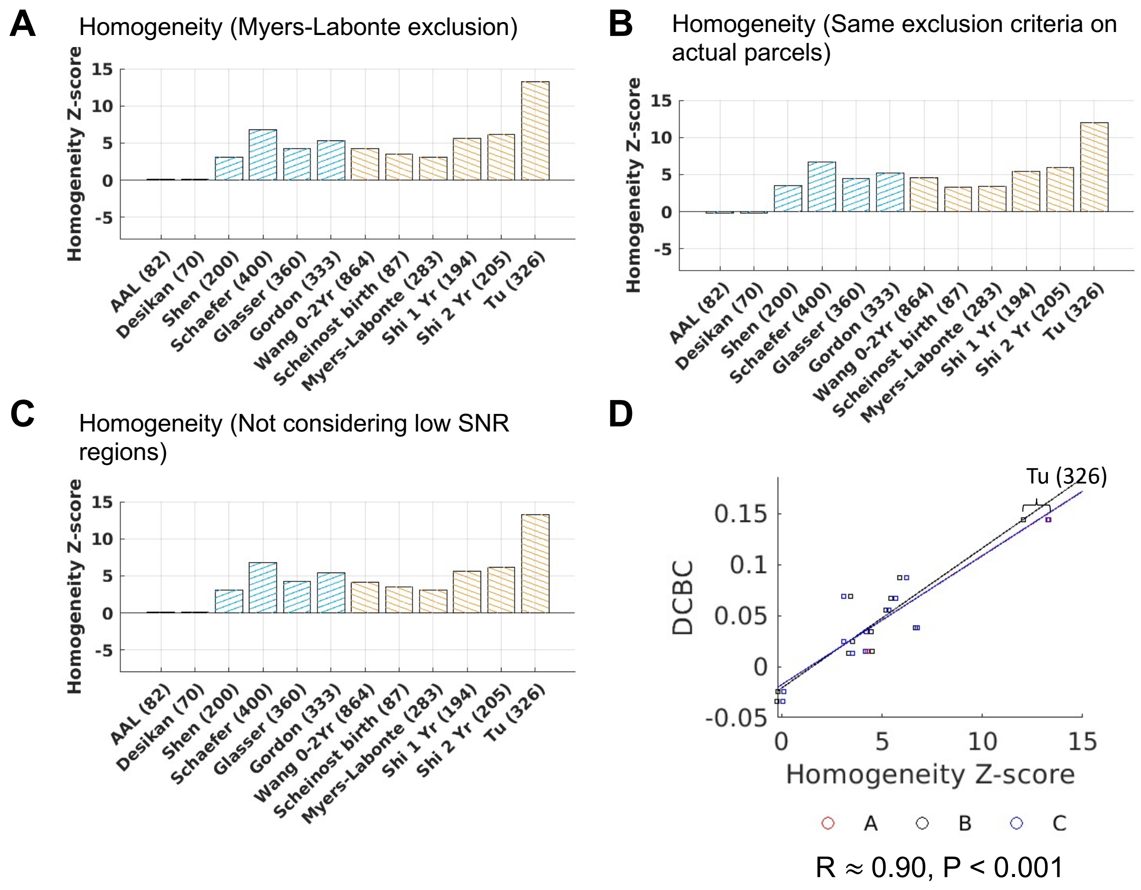


**Supplementary Figure 12.** *Functional connectivity homogeneity Z-scores using adult and early childhood parcellations on group-average eLABE (Y2) data (average homogeneity calculated using parcel-size weights).* A) Homogeneity Z-score (Myers-Labonte exclusion). B) Homogeneity Z-score (Same exclusion criteria on actual parcels). C) Homogeneity Z-score (Not considering low SNR regions). D) DCBC against homogeneity Z-score.

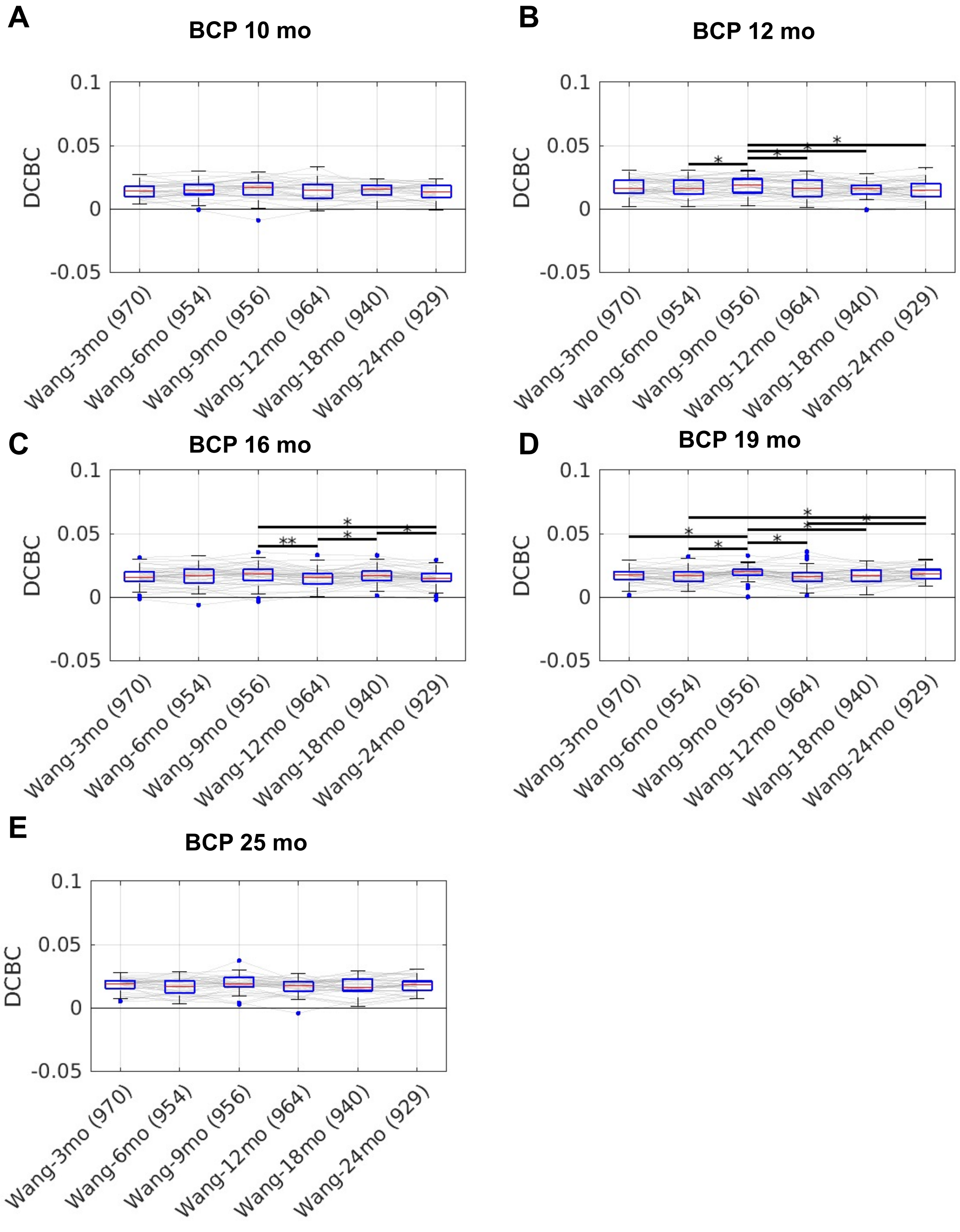


**Supplementary Figure 13.** *Distance-controlled boundary coefficient (DCBC) cluster validity for different versions of Wang parcellation.* *p<.05, **p<.01. Paired t-test FDR-corrected.

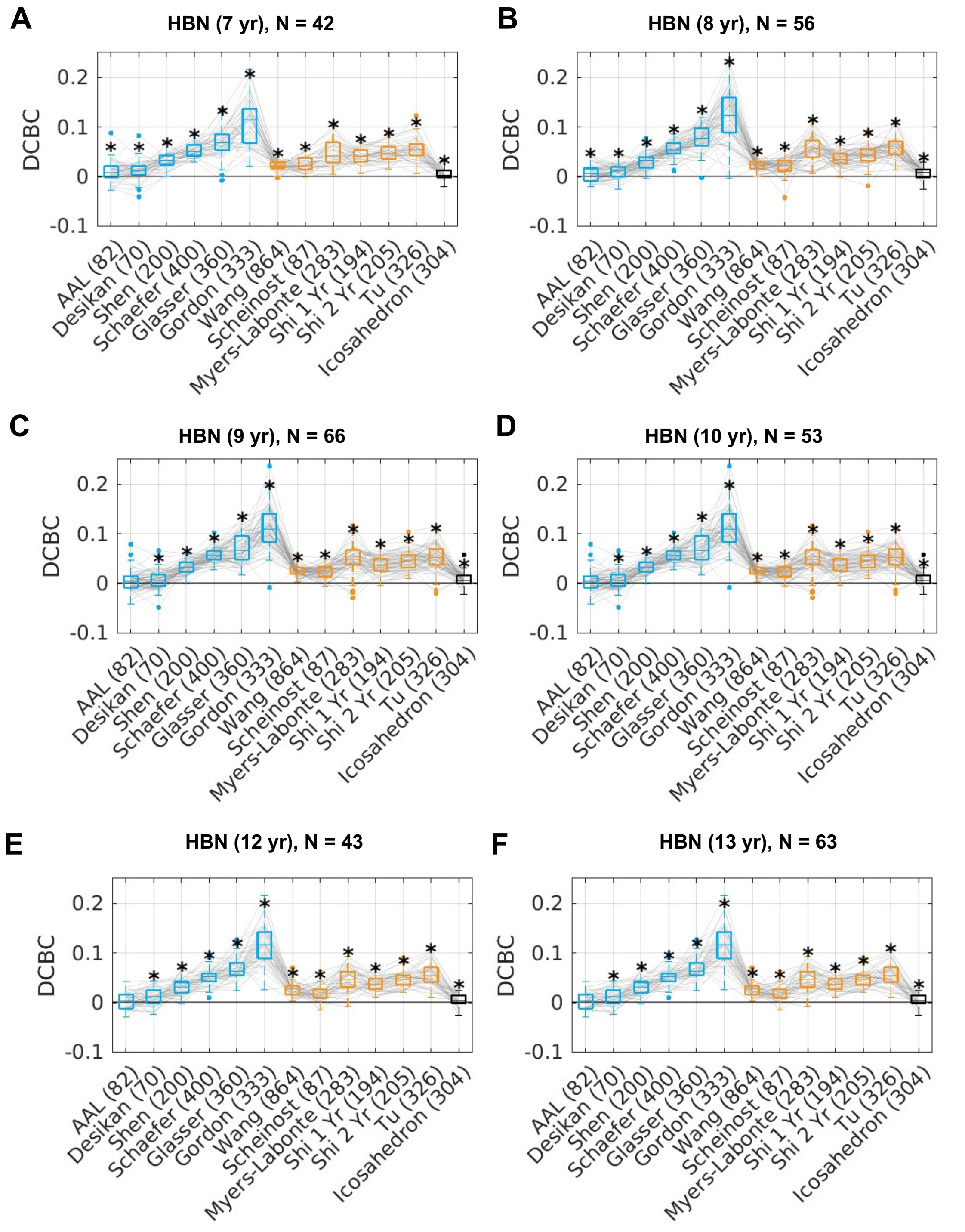


**Supplementary Figure 14.** *Distance-controlled boundary coefficient (DCBC) cluster validity for other HBN age groups*. * p<.05 after FDR-correction.

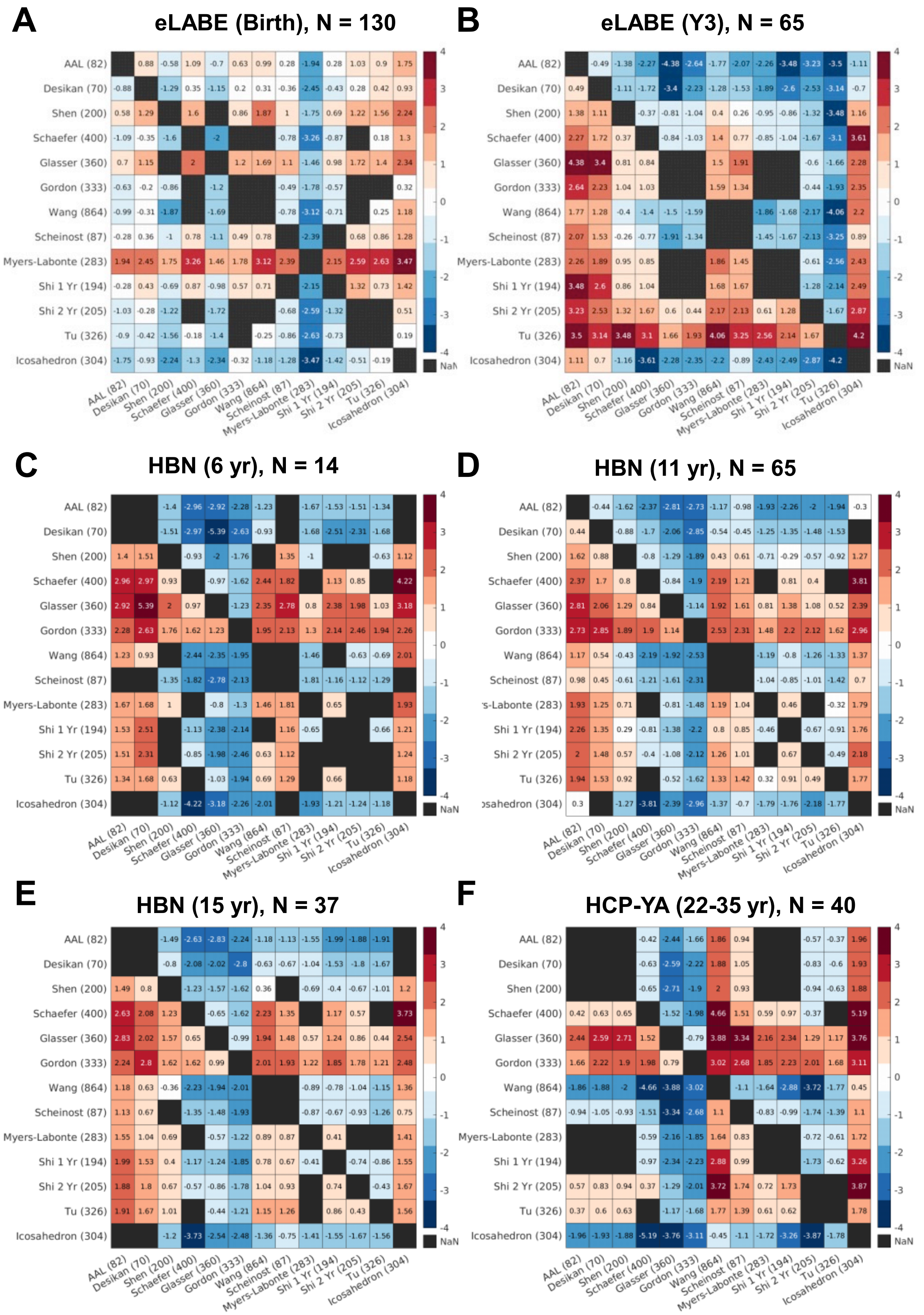

**Supplementary Figure 15.** *Effect size (Cohen's d for paired samples) of the difference in cluster validity between parcellations across datasets.* Each block shows the comparison between the parcellation in each row against the parcellation in each column. Non-significant differences were displayed as black blocks (NaN). A) eLABE (Birth). B) eLABE (Y3). C) HBN (6 yr). D) HBN (11 yr). E) HBN (15 yr). F) HCP-YA.

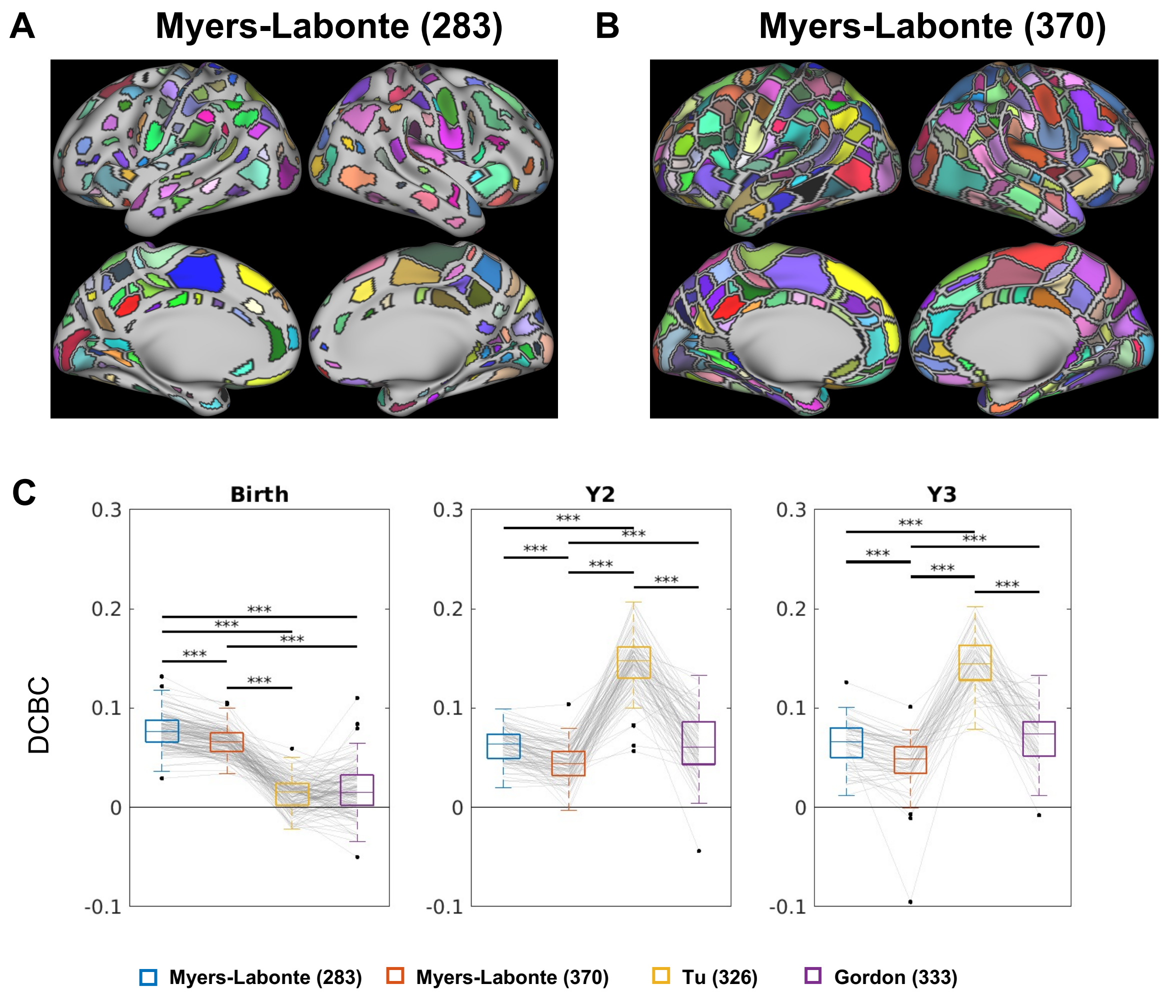


**Supplementary Figure 16.** *Distance-controlled boundary coefficient (DCBC) cluster validity for different versions of area parcellations from Myers et al. (2024) Cerebral Cortex*. A) Myers-Labonte (283). B) Myers-Labonte (370). C) The measure was evaluated on the different age groups of the eLABE dataset (birth to 3 years). The Birth data was a held-out subset not used in the development of Myers birth (283) or Myers birth (370) parcellations but the Y2 data was the same data used in the development of Tu (326) parcellation. *** p<.001. FDR-corrected for four paired t-tests.

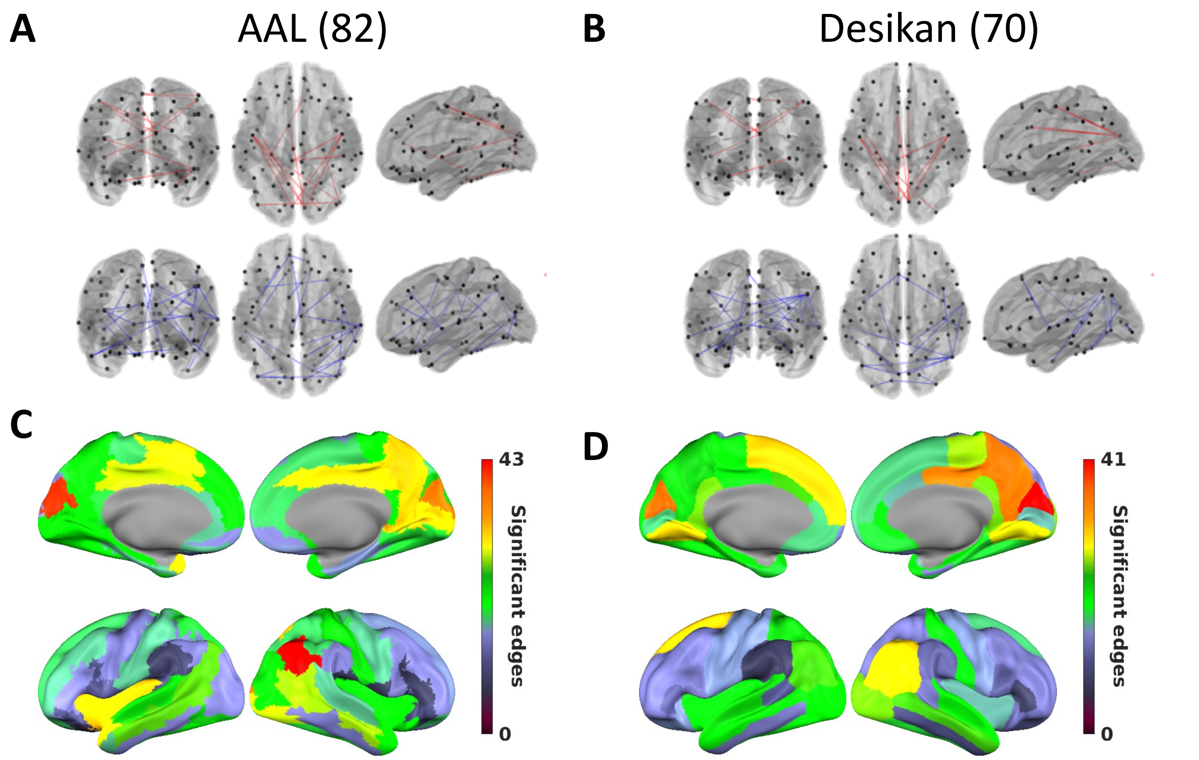


**Supplementary Figure 17.** *Predicting behavioral phenotypes from the connectomes using adult and early childhood parcellations (same as Figure 6 but for area parcellations: AAL (82) and Desikan (70)).* A/B/E/F: Top 5% positive edges in age-FC correlation magnitude (red) and top 5% negative edges in age-FC correlation magnitude (blue).

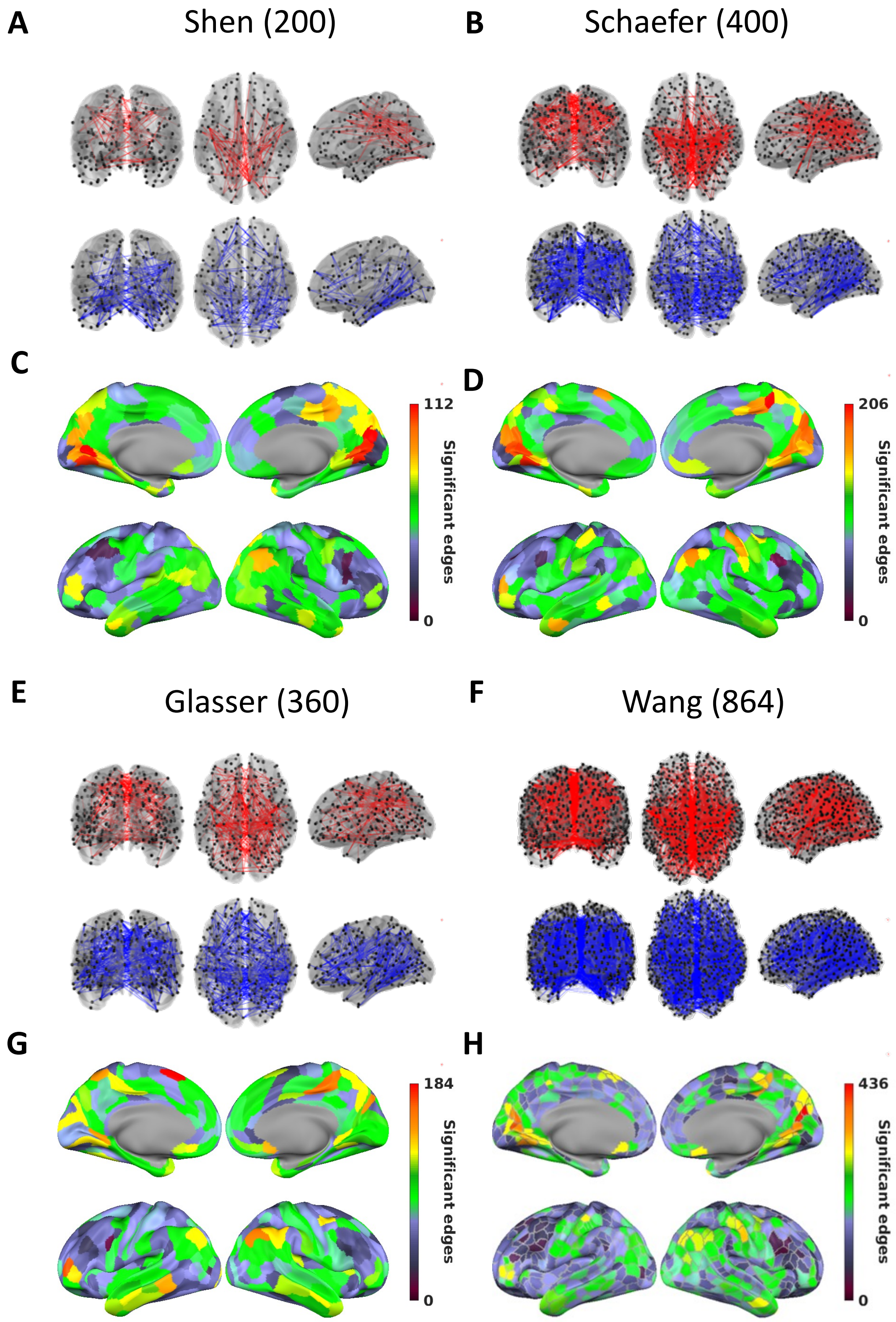


**Supplementary Figure 18.** *Predicting behavioral phenotypes from the connectomes using adult and early childhood parcellations* (same as Figure 6 but for area parcellations: Shen (200), Schaefer (400), Glasser (360) and Wang (864)). A/B/E/F: Top 5% positive edges in age-FC correlation magnitude (red) and top 5% negative edges in age-FC correlation magnitude (blue).

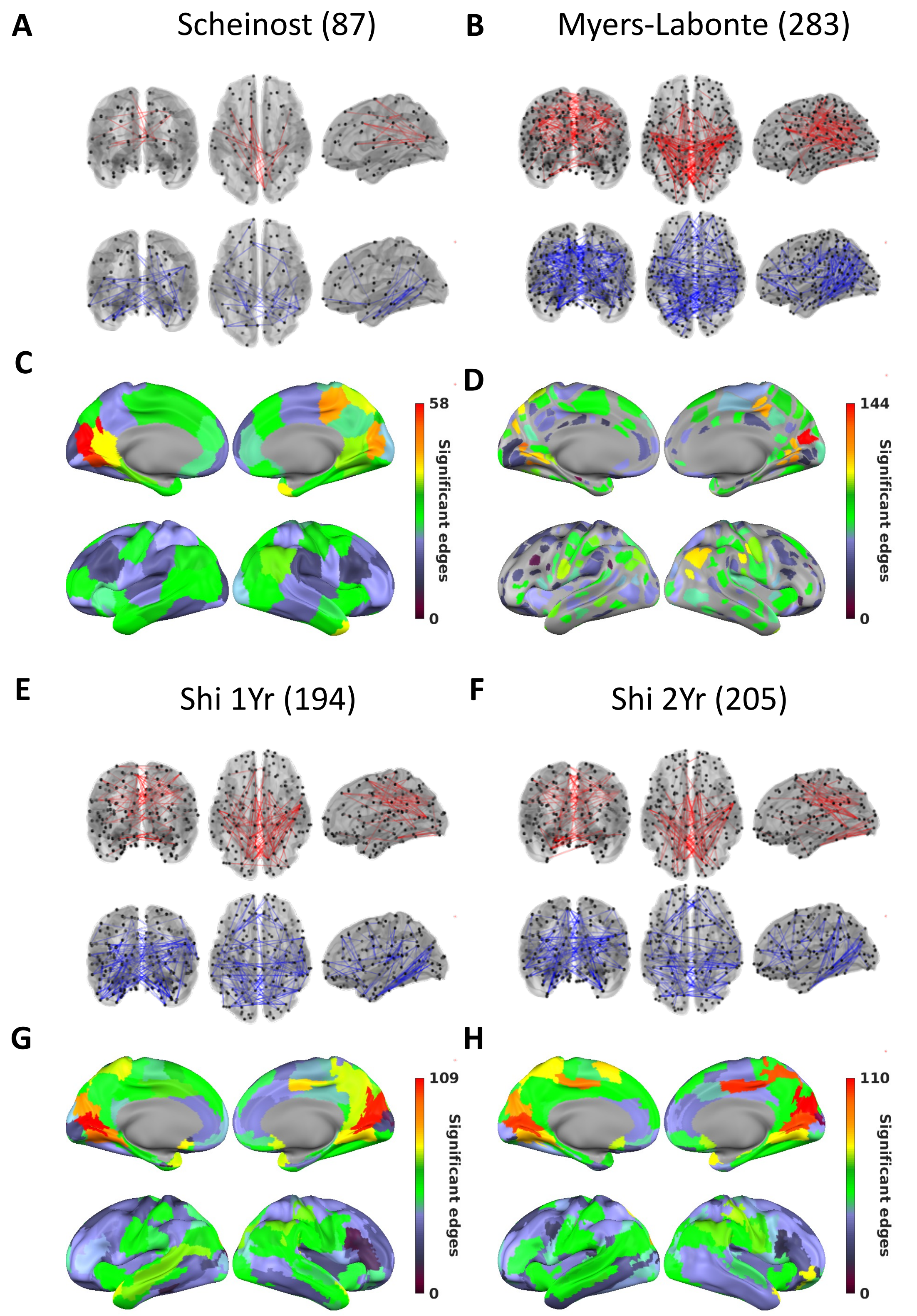


**Supplementary Figure 19.** *Predicting behavioral phenotypes from the connectomes using adult and early childhood parcellations* (same as Figure 6 but for area parcellations: Scheinost (87), Myers-Labonte (283), Shi 1 Yr (194) and Shi 2Yr (205)). A/B/E/F: Top 5% positive edges in age-FC correlation magnitude (red) and top 5% negative edges in age-FC correlation magnitude (blue).

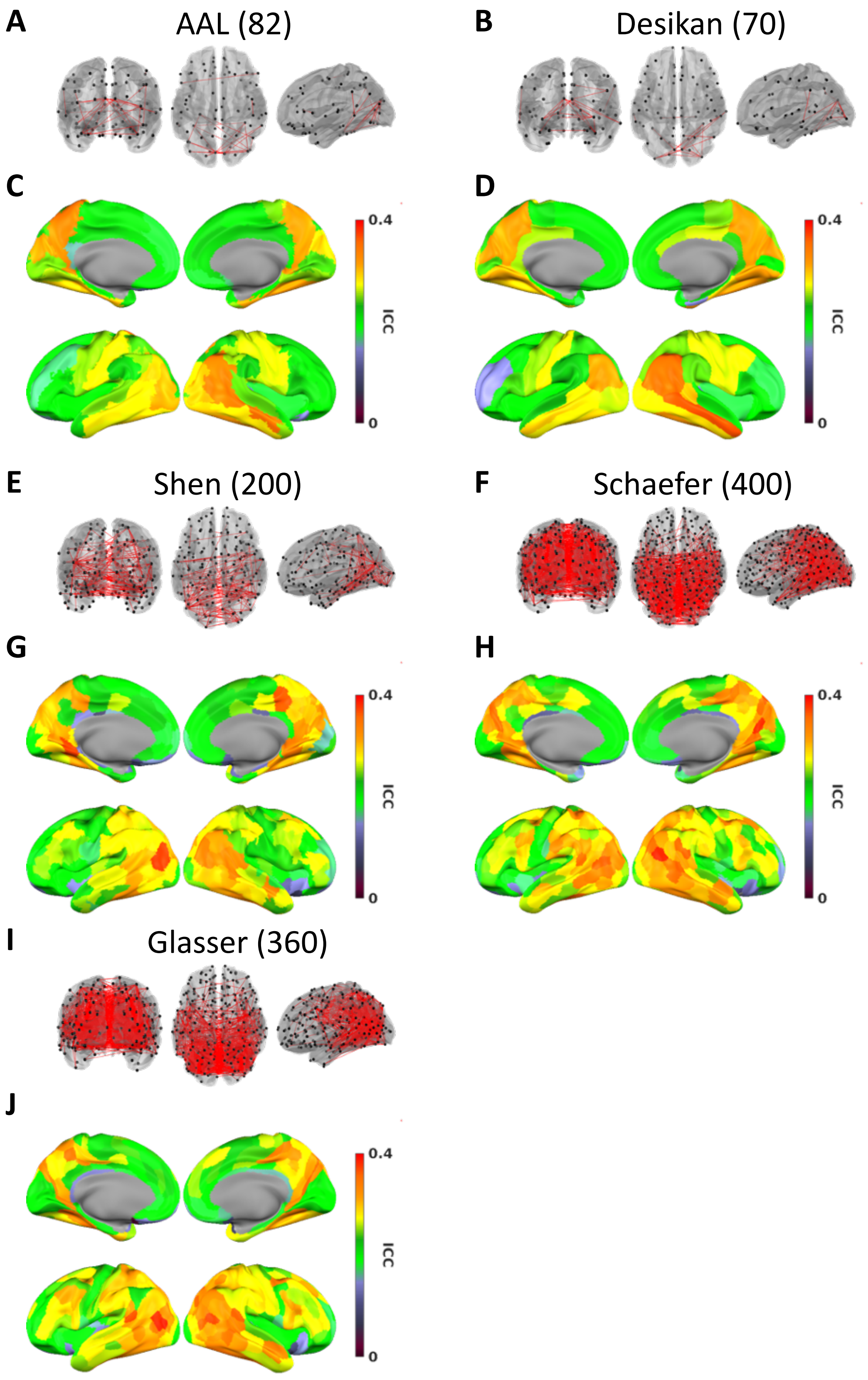


**Supplementary Figure 20.** *Test-retest reliability from the connectomes using adult parcellations (same as Figure 7 but for other adult area parcellations)*. A/B/E/F/I: The edges with a "good" reliability (ICC = 0.60-0.75). C/D/G/H/J: Mean edge test-retest reliability for edges connected to areas.

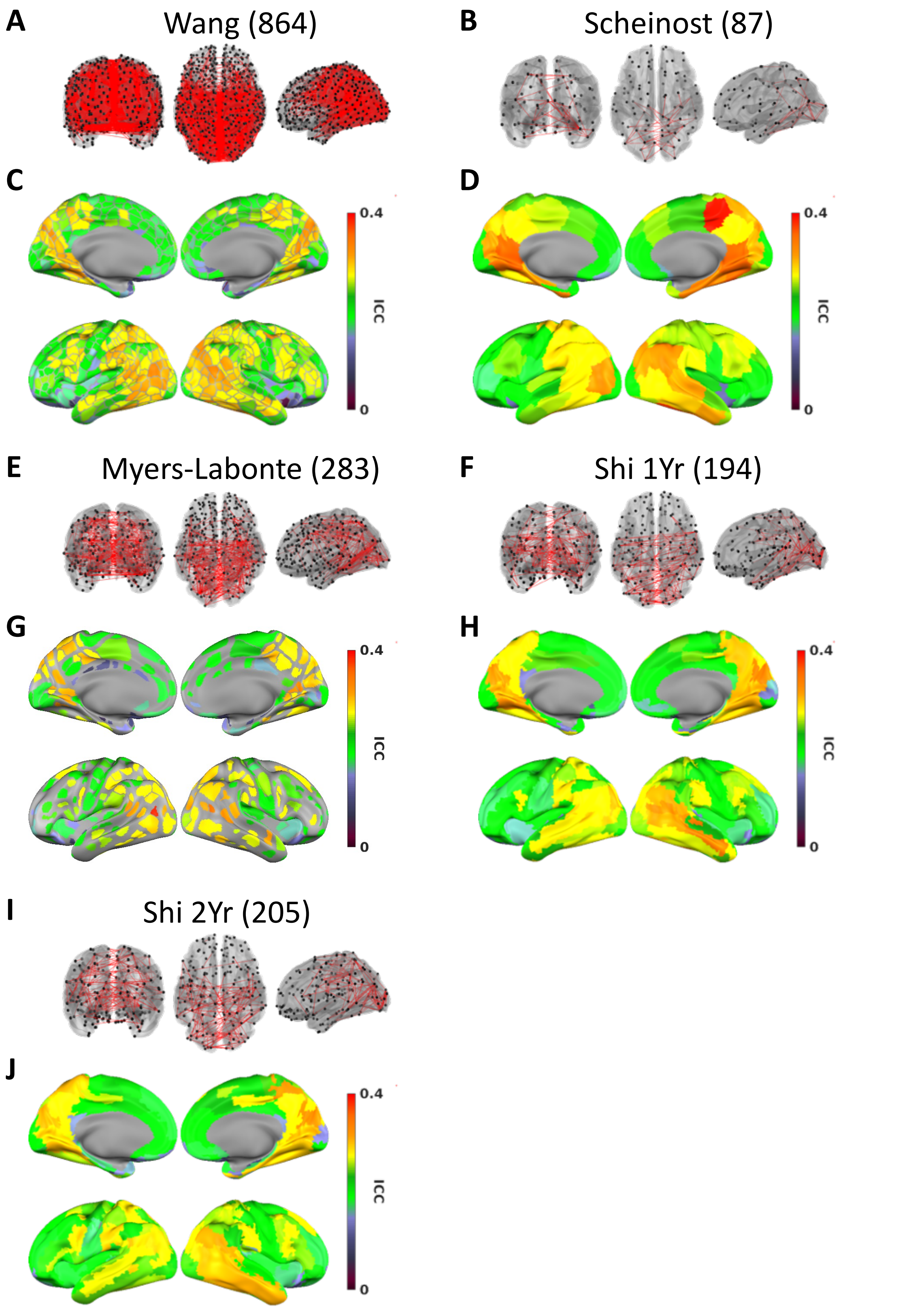


**Supplementary Figure 21.** *Test-retest reliability from the connectomes using early childhood parcellations (same as Figure 7 but for other early childhood area parcellations)*. A/B/E/F/I: The edges with a "good" reliability (ICC = 0.60-0.75). C/D/G/H/J: Mean edge test-retest reliability for edges connected to areas.

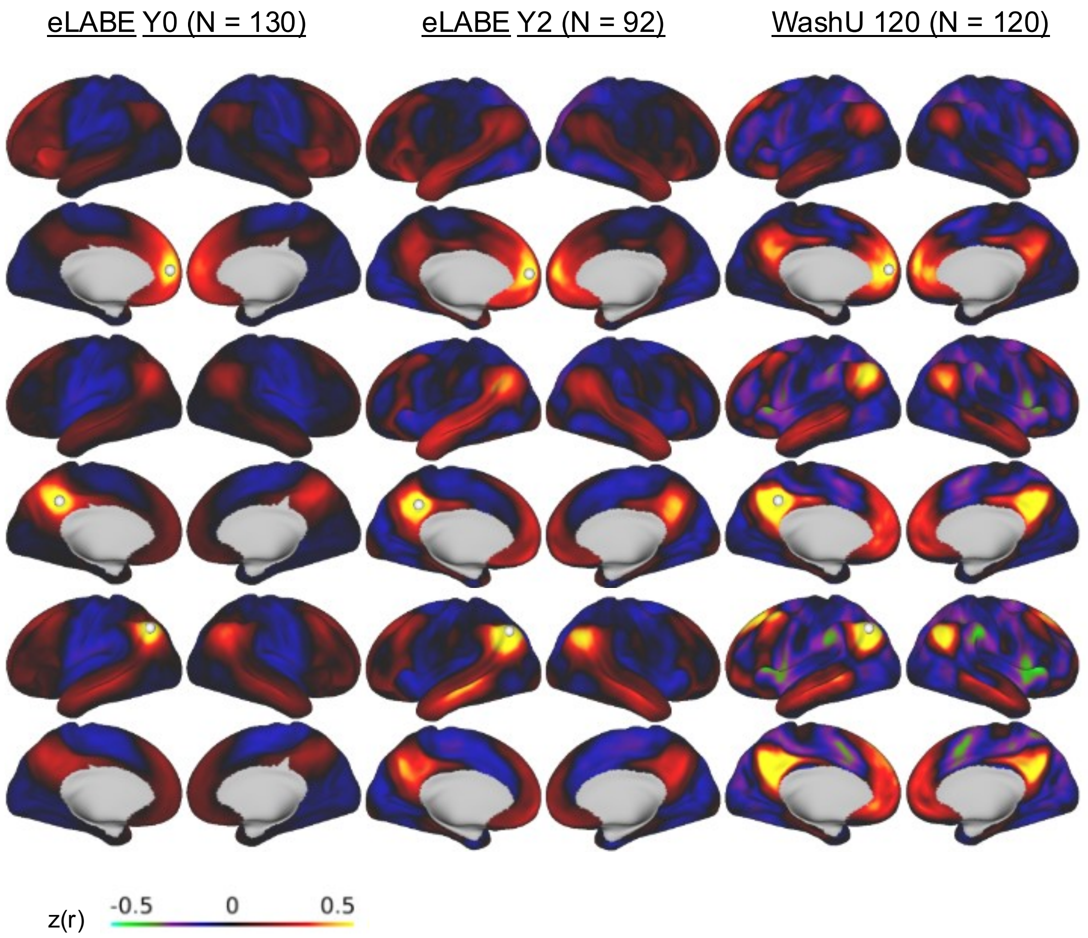


**Supplementary Figure 22.** *Raw connectivity seed maps (color scales with z(r)) for neonates, toddlers and adults from anterior (Top), posterior (Middle) and temporal (Bottom) default mode system.*

| Supplementary Table 1 | | | |
|---|---|---|---|
| | Cohort | BCP | eLABE Y2 |
| **Acquistion** | Location | University of Minnesota | Washington University in St. Louis |
| | Scanners | Siemens Prisma 3T Scanner | Siemens Prisma 3T Scanner |
| | Headcoil | 32-channel | 64-channel |
| | Sequence type | Gradient-echo EPI | Gradient-echo EPI |
| | Resolution (BOLD) | 2mm isotropic | 2mm isotropic |
| | Phase encoding direction | AP+PA | AP |
| | TR (s) | UMN - 0.72 (N = 70), 0.8 (N = 107) | 0.8 |
| | TE (ms) | 37 | 37 |
| | Resolution (T1) | 0.8 mm | 0.8mm |
| | Multi-band factor | 8 | 8 |
| | State | Natural Sleep | Natural Sleep |
| | Total acquistion time | Mean 1253 frames (13.1 min) | 20.0 min (range 11.2-42.5 min) |
| | Frames per BOLD run | 420 | 420 |
| | BOLD runs | 2-4 | 1-8 |
| **Processing** | Processing pipeline version | DCAN-Infant (0.0.9) | toddler EPI (BOLD) preprocessing pipeline using the 4dfp tool suite |
| | Distortion correction | ANTs SyN registration | FSL topup |
| | Bias field correction | N4 method | FSL fast |
| | Denoising | Respiratory filter, demean, detrend, 24 parameters nuisance regression, remove FD>0.3mm to apply bandpass filtering (0.008-0.09 Hz) then interpolate the missing frames | Nuisance waveforms were regressed out of the time- series including retrospective motion correction from the 24-Friston parameters, gray matter global signal, and regions of non-interest and their first derivative (white matter, ventricular, and extra-axial CSF). The data were then bandpass filtered in the |

| | | | |
|---|---|---|---|
| | | | range of 0.005 to 0.1 Hz to eliminate non-BOLD frequencies |
| | Scrubbing threshold | filtered FD<0.2mm, outlier (across-vertex STD on low FD frames >3MAD of the median of all frames) | filtered FD<0.2mm and at least 3 consecutive low-motion frames |
| | Tissue Segmentation | ANTs Joint Label Fusion was performed using a set of ALBERT atlases with segmentations generated and manually corrected by DCAN for ages 0-5 months old. For older ages, the atlases used for JLF were a set of 10 ABCD subjects for which we generated segmentations. Manual curation of tissue segmentation was performed where necessary. | FreeSurfer default recon-all processing pipeline. Manual curation of tissue segmentation was performed where necessary. |
| | Surface reconstruction & registration | Modified FreeSurfer reconstruction (no hires, aseg from JLF, adjusted class means of tissue to fit T1w contrasts), registered to fs_LR32k using spherical registration (with MSMsulc). | BOLD-> individualT1-> cohort-specific atlas ->711-2B Talairach Atlas (linear with 4dfp_tools). T1-weighted MRI images were registered and resampled to an age-specific atlas-representative target. This volumetric timeseries was then registered to an adult atlas target (711–2B), then registered to fs_LR32k using spherical registration. |
| | BOLD data geodesic smoothing | σ = 2.55 mm | σ = 2.25 mm |
| **Quality Control** | | | |
| | | BrainSwipes crowdsource ratings with average aggregated passing rate >75% for anatomical or functional images + manual screening. | The registration of the T1 - weighted image, atlas target, and rs-fMRI timeseries were manually inspected by experienced raters to ensure accuracy of individual processing result. |
| | References | Kardan 2022 DCN | Kaplan 2021 Neuroimage |
| | Data availability | https://nda.nih.gov/edit_collection.html?id=2848 | https://eedp.wustl.edu/research/elabe-study/ |

| **Supplementary Table 2** | | | |
|---|---|---|---|
| | Cohort | eLABE Birth | WashU 120 |
| **Demographics** | Age | 41.6 weeks (SD = 1.3 weeks) | Mean 25 yr, range 19-32 yr |
| | Subjects | 131 | 120 |
| | Sex (M/F) | 78/53 | 60/60 |
| | Race | 48 white, 81 black, 3 American Indian or Alaska native | N/A |

| | | | |
|---|---|---|---|
| **Acquistion** | Location | Washington University in St. Louis | Washington University in St. Louis |
| | Scanners | Siemens Prisma 3T Scanner | Siemens 3T Trio Tim |
| | Headcoil | 64-channel | 12-channel |
| | Sequence type | Gradient-echo EPI | Gradient-echo EPI |
| | Resolution (BOLD) | 2mm isotropic | 4 mm isotropic |
| | Phase encoding direction | AP+PA | AP |
| | TR (s) | 0.8 | 2.5 |
| | TE (ms) | 37 | 27 |
| | Resolution (T1) | 0.8mm | 1 mm |
| | Multi-band factor | 8 | N/A |
| | State | Natural Sleep | Fixation on crosshair |
| | Total acquistion time | 22.8 min (range 16.8-44.8 min) | 14.0 min (range 7.7-30.4 min) |
| | Frames per BOLD run | 420 | 120 |
| | BOLD runs | 2-9 | 2 |
| **Processing** | Processing pipeline version | neonate EPI (BOLD) preprocessing pipeline using the 4dfp tool suite | adult EPI (BOLD) preprocessing pipeline using the 4dfp tool suite |
| | Distortion correction | FSL topup | None |
| | Bias field correction | FSL fast | None |
| | Denoising | Demean, detrend, 24 parameters nuisance regression (using gray matter signal as whole-brain signal), bandpass filter (0.005-0.1 Hz) | Demeaning and detrending, , multiple regression including: whole-brain, ventricular and white matter signals, and motion regressors derived by Volterra expansion (Friston et al. 1996), and a band-pass filter (0.009 Hz < f < 0.08Hz). |
| | Scrubbing threshold | FD<0.25 mm and at least 3 consecutive low-motion frames | FD<0.2mm, at least 5 consecutive low-motion frames |
| | Tissue Segmentation | Segmentation was performed with the M-CRIB atlas. Manual curation of tissue segmentation was performed where necessary. | FreeSurfer's default recon-all processing pipeline (version 5.0) |
| | Surface reconstruction & registration | BOLD-> individualT2-> cohort-specific atlas ->711-2B Talairach Atlas (linear with 4dfp_tools). Subject T2 linearly transformed to adult Talairach space and reconstructed subject-specific surface with M-CRIB atlas, then | FreeSurfer's default recon-all processing pipeline (version 5.0) |

| | | | |
|---|---|---|---|
| | | registered to fs_LR32k using spherical registration. | |
| | BOLD data geodesic smoothing | σ = 1mm before noise removal and filtering,σ = 2.25 mm after | σ = 2.55 mm |
| **Quality Control** | | | |
| | | The registration of the T2 -weighted image, atlas target, and rs-fMRI timeseries were manually inspected by experienced raters to ensure accuracy of individual processing result. | N/A |
| | References | Sylvester 2022 Cerebral Cortex | Gordon 2016 Cerebral Cortex |
| | Data availability | https://eedp.wustl.edu/research/elabe-study/ | https://legacy.openfmri.org/dataset/ds000243/ |
